# Point mutations in the FimH lectin domain: predicted folding stability versus conformational dynamics for R60P, Q133K and Y137A

**DOI:** 10.64898/2026.11.10.765513

**Authors:** Jose Angel Navas-Passos, Adan Pereira-Tapia, Carlos Velez-Duncan, Antistio Alviz-Amador

## Abstract

**Background:** The adhesin FimH attaches uropathogenic *Escherichia coli* to the urinary epithelium through a force-regulated catch-bond that single lectin-domain substitutions can modulate. How such substitutions reshape the dynamics of the isolated lectin domain, and whether stability predictors anticipate them, is unclear.

**Methods:** We compared the wild-type (WT) lectin domain (PDB 4XO8, apo) with R60P, Y137A and Q133K, using three 500 ns molecular dynamics (MD) replicates per system (6 microseconds total) and three folding-stability predictors (FoldX, a Cartesian-ddG-type Rosetta protocol, ERIS-2). The replicate was the unit of analysis (n = 3), with 95% confidence intervals and a WT replicate-to-replicate control.

**Results:** FoldX and ERIS-2 ranked the variants consistently, R60P most destabilising and Q133K neutral; the Rosetta value for Q133K was anomalously high. All systems stayed folded. R60P gave the only reproduced local change, loss of beta-strand structure at residues 60-61 and of three local hydrogen bonds. Y137A did not bring Thr158 into the pocket, and the Q133K Gln133-Asp54 polar contact became a Lys133-Asp54 salt bridge. In the principal-component landscape, departures from the WT region came from individual replicates (Y137A replicate 1; Q133K replicates 2-3), not a systematic shift.

**Conclusions:** A substitution perturbs local structure most clearly when it breaks a beta-strand (R60P), and the stability predictions and dynamics together give a consistent ranking of the variants, including the first dynamic characterisation of Q133K. The results characterise the apo lectin domain and give a reproducible basis for interpreting FimH mutations, not for inferring affinity.

## 1. Introduction

Urinary tract infections (UTIs) are among the most common bacterial infections worldwide, and uropathogenic Escherichia coli (UPEC) is their most frequent cause in uncomplicated and complicated disease (Mancuso et al., 2023; Whelan et al., 2023). An estimated 40-50% of women experience at least one symptomatic UTI during their lifetime (Hatton et al., 2021), and the clinical burden is compounded by antimicrobial resistance: in 2019, bacterial resistance in UTIs was estimated to be attributable to roughly 64,890 deaths and associated with about 260,000, with E. coli and Klebsiella pneumoniae responsible for more than half and with high resistance to fluoroquinolones and third-generation cephalosporins (Li et al., 2022). This motivates strategies that do not rely on killing the bacterium, such as blocking adhesion to the urothelium (Hatton et al., 2021; Sarshar et al., 2020).

Adhesion of UPEC to the bladder is mediated by FimH, an adhesin at the tip of type 1 pili that recognises mannose-rich glycans on the urinary epithelium (Sarshar et al., 2020). FimH comprises a lectin domain, which houses the mannose-binding pocket, and a pilin domain that anchors it to the pilus (Figure 1). Its mechanical behaviour is unusual: adhesion increases under shear stress, allowing the bacterium to resist urine flow (Thomas et al., 2002), a behaviour explained by a catch-bond in which ligand affinity rises when force is applied (Sauer et al., 2016).

**Figure 1.**
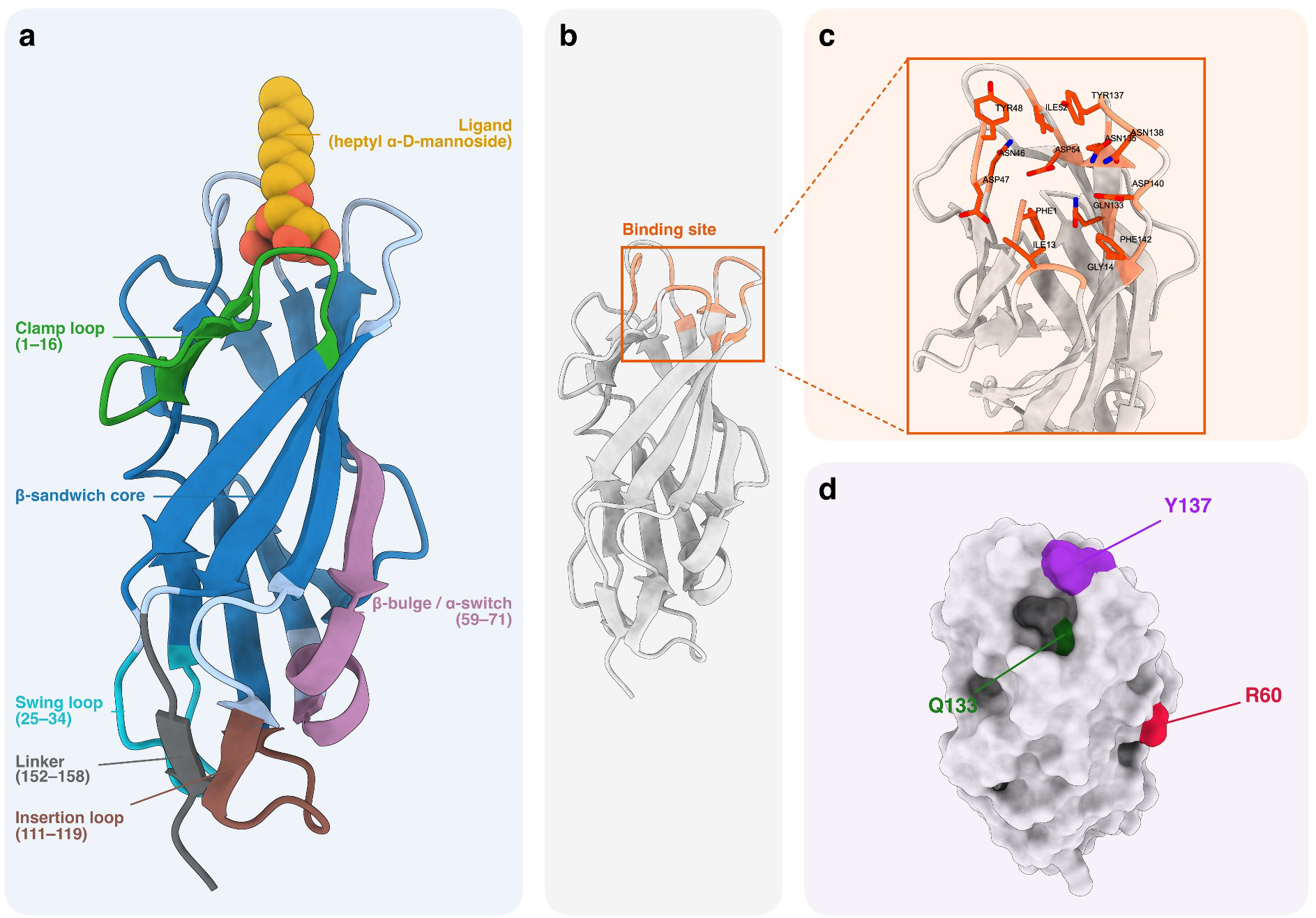
Structure of the FimH lectin domain (author-prepared in ChimeraX). (a) Monomeric lectin domain (PDB 4XO8, chain A) with heptyl α-D-mannoside (spheres); structural regions coloured and labelled per Languin-Cattoën et al. (2023): clamp loop (1-16), swing loop (25-34), β-bulge/α-switch (59-71), insertion loop (111-119), linker (152-158) and the β-sandwich core. (b) Same orientation without ligand; residues within 4 Angstrom of the ligand are highlighted as the binding site. (c) Close-up of the binding site with labelled residues. (d) Molecular surface with the mutated positions R60, Q133 and Y137 highlighted.

The dominant structural model proposes that, at rest, the lectin domain is coupled to the pilin domain in a low-affinity conformation; tension separates the domains and twists the central beta-sheet in a finger-trap-like manner, driving the pocket to a high-affinity state (Le Trong et al., 2010). MD simulations indicate that this beta-sheet behaves as a soft spring whose twist correlates with fluctuations of the interdomain region and the binding site, and that the pocket is more rigid and excludes water better in the high-affinity state (Interlandi & Thomas, 2016). Yet the two states show nearly superimposable pocket geometries, and enhanced sampling shows that apparent affinity depends on binding-site populations the crystal does not reveal (Languin-Cattoën et al., 2023).

Three lectin-domain substitutions are of particular interest. Arg60-to-Pro (R60P) stabilises the isolated lectin domain in the low-affinity conformation and has been proposed as a minimal model of FimH, with simulations reporting larger backbone fluctuations in the pocket and interdomain region than in the high-affinity lectin domain (Liu et al., 2021). Tyr137-to-Ala (Y137A) is the most detrimental of the tyrosine-gate mutations for affinity and specificity: in the absence of ligand, the C-terminal residue Thr158 intrudes into the pocket, and relaxation of the backbone dihedrals at position 137 couples to Tyr48 through Ile52 (Rabbani et al., 2017). Finally, Gln133 forms a hydrogen bond with the mannose O3 that simulations find stable only in the high-affinity state (Interlandi & Thomas, 2016); we found no study characterising Gln133-to-Lys (Q133K). Natural mutations also tune allostery: the FimH30 variant of the multidrug-resistant ST131 lineage carries R166H, which weakens interdomain interactions and is associated with greater adhesion (Qin et al., 2022).

In parallel, the thermodynamic stability of the native fold frequently determines the effect of a substitution: positions of pathogenic mutations are more prone to destabilise the fold than neutral ones (Redler et al., 2016). However, ΔΔG predictors have limited accuracy and are not always consistent between the forward and reverse mutation (Thiltgen & Goldstein, 2012); for FoldX, the uncertainty of an individual prediction can reach ±2.9 kcal/mol and improves when MD is incorporated (Sapozhnikov et al., 2023). Previous work has combined ΔΔG and MD to interpret point substitutions (Williams et al., 2017; Zhang et al., 2021), whereas single-trajectory comparisons (Mir et al., 2024) cannot quantify inter-replicate variability.

To our knowledge, the predicted stability and the dynamics of the apo FimH lectin domain have not been compared under identical conditions for these three substitutions against the wild type, using independent replicates that estimate sampling variability. This comparison indicates how sensitive the pocket and allosteric region are to a single substitution, informing antagonist and variant design, and provides a reproducible framework, with explicit uncertainty thresholds, for FimH mutations.

The general objective was to evaluate how R60P, Y137A and Q133K modify the predicted stability and conformational dynamics of the apo FimH lectin domain relative to the wild type. Three hypotheses were pre-specified before analysing the trajectories. H1: R60P alters local flexibility and pocket geometry. H2: Y137A changes tyrosine-gate dynamics and the position of Thr158. H3: Q133K establishes new interactions with Asp47/Asp54/Asp140 and the N-terminal amine. Null outcomes are reported in the same detail as positive ones.

## 2. Methods

### 2.1 Study design and scope

A comparative computational study was carried out with four systems: the wild-type lectin domain (WT) and three single-substitution variants (R60P, Y137A, Q133K), each simulated in three independent 500 ns replicates (12 trajectories, 6 microseconds). The study is restricted to the isolated lectin domain in the apo state: it includes neither the pilin domain, nor ligand, nor applied force. Consequently, the results describe folding stability and local dynamics of the lectin domain, not mannose affinity or adhesion. Because FimH dynamics are slow, 500 ns may be not enough to observe complete transitions between affinity states, so the analysis focuses on changes in flexibility and local organisation of the pocket.

### 2.2 Starting structure and mutagenesis

The crystal structure of the E. coli FimH lectin domain was used (PDB 4XO8, 1.70 Angstrom resolution). With Biopython (Cock et al., 2009), chain A (residues 1-158) was retained and the co-crystallised heptyl mannoside and all crystallographic waters were removed to obtain an apo structure. Identity at positions 60, 133 and 137 (Arg, Gln, Tyr) was verified before mutation, and the 4XO8 numbering is used throughout. Three independent point mutants were generated with PDBFixer (Eastman et al., 2017; applyMutations), and the unmutated structure was kept as WT. Hydrogens were added at pH 7.4 identically for the four systems; His45 was assigned as HIE and the Cys3-Cys44 disulfide was preserved. In R60P, an open proline ring produced by PDBFixer was rebuilt and its geometry was verified in the trajectories (Supplementary Material). Molecular structures were rendered with UCSF ChimeraX (Pettersen et al., 2021).

### 2.3 System construction, equilibration and production

Systems were built with tleap (Case et al., 2023) using the ff14SB force field (Maier et al., 2015) and TIP3P water (Jorgensen et al., 1983) in a truncated octahedral box with a 10.0 Angstrom minimum solute-to-boundary distance. Each system was neutralised and Na+/Cl- pairs were added to approximately 150 mM, the number of pairs being round (0.150 × N_water / 55.5). The alkali-halide (Na+, Cl-) parameters were those of Joung and Cheatham (2008), within the standard TIP3P ion set whose remaining monovalent parameters follow Li et al. (2015). Amber naming conventions were applied (His45 as HIE; Cys3 and Cys44 as CYX for the disulfide; the N-terminal hydrogen as H1). The resulting atom counts, ion numbers, salt concentrations, net charges and initial box sizes are given in Table S2; the R60P and Q133K substitutions change the net charge by −1 and +1, so differences for these variants cannot be attributed with certainty to the structural substitution rather than to charge. For each replicate, ions were repositioned with cpptraj randomizeions (Roe & Cheatham, 2013) at least 6.0 Angstrom from the solute and 4.0 Angstrom from one another, using a distinct seed.

Equilibration used a 2 fs time step, an 8.0 Angstrom cutoff, SHAKE on bonds to hydrogen (Ryckaert et al., 1977), particle-mesh Ewald electrostatics (Darden et al., 1993) and harmonic position restraints on the heavy atoms of residues 1-158 referenced to the initial coordinates. It proceeded through nine stages — four restrained minimisations, a 50 ps restrained NVT stage at 300 K, and four 100 ps NPT stages at 1 atm that progressively released the restraints (Table S3) — with a Berendsen thermostat and barostat (Berendsen et al., 1984; τ_T = 0.5 ps, τ_P = 1.0 ps). Hydrogen-mass repartitioning (Hopkins et al., 2015; H = 3.024 u), generated after equilibration and used only in production, allowed a 4 fs time step. Production was run with pmemd.cuda of Amber 26 at 300 K using a Langevin thermostat (γ = 2.0 ps⁻¹) and a Monte Carlo barostat (Åqvist et al., 2004) at 1 atm, an 8.0 Angstrom cutoff, SHAKE, and removal of centre-of-mass motion every 1000 steps; coordinates were saved every 10 ps. Each replicate was run for 500 ns as ten chained 50 ns segments with continuous velocities and time, with a per-replicate random seed (*SEED = 10000 × (system index + 1) + replicate; production ig = SEED + 500000*). The three replicates of a system therefore share the same starting conformation and box and differ only in initial velocities and ion placement. Software versions are listed in the Supplementary Material.

### 2.4 ΔΔG prediction

Folding stability was estimated with three predictors that use different units and, in one case, a different input structure, so their absolute values are not comparable and only the rank order is interpreted (Figure 4g, Table 1). FoldX 5.1 (Schymkowitz et al., 2005) was run with RepairPDB on the repaired WT structure derived from 4XO8, and each substitution was modelled with BuildModel in five runs (mean ± SD; positive = destabilising, kcal/mol). A Cartesian-ddG-type protocol was run in PyRosetta with the ref2015_cart energy function (Park et al., 2016): Cartesian FastRelax, repacking of neighbours within 8 Angstrom and minimisation, over three independently relaxed structures, in Rosetta energy units (REU); this is a custom approximation, not the official cartesian_ddg binary, and REU are not converted to kcal/mol. ERIS-2 was run in its empirical (“classic”) mode, reporting ΔΔG in kcal/mol. The input structure for the classic ERIS-2 run (Figure 4g, Table 1) is being confirmed against the FoldX input, and these values are therefore used only for the rank order (see the footnote to Table 1). A neural-mode ERIS-2 run returned neutral values for all three variants and is reported in the Supplementary Material only. Predictions are interpreted relative to the FoldX uncertainty of up to ±2.9 kcal/mol (Sapozhnikov et al., 2023).

**Table 1.** Predicted folding-stability change (ΔΔG) per mutation and method (shown in Figure 4g). Units differ and are not interchangeable; n denotes internal runs, not independent biological replicates. The Rosetta protocol is a custom Cartesian-ddG-type approximation (REU), not the official binary. ERIS-2 values are the empirical (“classic”) mode. No statistical test is applied between methods; positive values indicate destabilisation.

| **Mutation** | **FoldX (kcal/mol, mean ± SD, n = 5)** | **Rosetta (REU, mean ± SD, n = 3)** | **ERIS-2 classic (kcal/mol) ‡** |
|---|---|---|---|
| **R60P** | 3.84 ± 0.19 | 17.97 ± 2.03 | 25.23 (Destabilizing) |
| **Y137A** | 2.01 ± 0.03 | 2.07 ± 0.02 | 8.82 (Destabilizing) |
| **Q133K** | 0.60 ± 0.28 | 13.39 ± 0.53 † | 0.21 (Neutral) |

*† Rosetta Q133K is discordant with FoldX, ERIS-2 and the MD and is treated as a probable artefact of the custom protocol for a charge-changing substitution. ‡ ERIS-2 classic values; the input structure for this run is being confirmed against the FoldX input, so these values are used only for the rank order. A neural-mode ERIS-2 run returned 0.01 / 0.15 / 0.46 kcal/mol (all neutral), reported in the Supplementary Material only.*

### 2.5 Trajectory analysis and two pipelines

Analyses used the 100-500 ns window (first 100 ns discarded) and two complementary pipelines. (i) Global metrics in Figure 2 — backbone (N, Cα, C) RMSD, per-residue Cα RMSF, radius of gyration and total SASA — were computed with cpptraj (Roe & Cheatham, 2013) after superposition on the backbone. (ii) Binding-pocket RMSD (Figure 3a-b) was computed from a separate series on the thirteen binding-site residues defined in the project methodology (Phe1, Ile13, Asn46, Asp47, Tyr48, Ile52, Asp54, Tyr95, residue 133, Asn135, residue 137, Asp140, Phe142). (iii) Essential dynamics (PCA, free-energy landscapes), difference dynamic cross-correlation maps (ΔDCCM), secondary structure (DSSP; Kabsch & Sander, 1983) and the hydrogen-bond/salt-bridge network (Figures 3c-g and 4) were computed on a covariance matrix and contact definitions common to the four systems. Because pipelines (i) and (ii-iii) differ, the same quantity can take slightly different absolute values between figures (for example, radius of gyration and total SASA). Three distinct operational definitions of the “pocket” appear in the figures and are kept separate: residues within 4 Angstrom of the ligand (Figure 1b-c), the thirteen binding-site residues above (Figure 3a-b), and an 8 Angstrom shell around each mutated site (Supplementary). Principal components were obtained from the Cα covariance of all systems pooled; free-energy landscapes were estimated as ΔG = -k_B T ln P at 300 K from the PC1-PC2 histogram, masked where sampling was sparse, and are shown as sampling estimates rather than converged thermodynamics.

**Figure 2.**
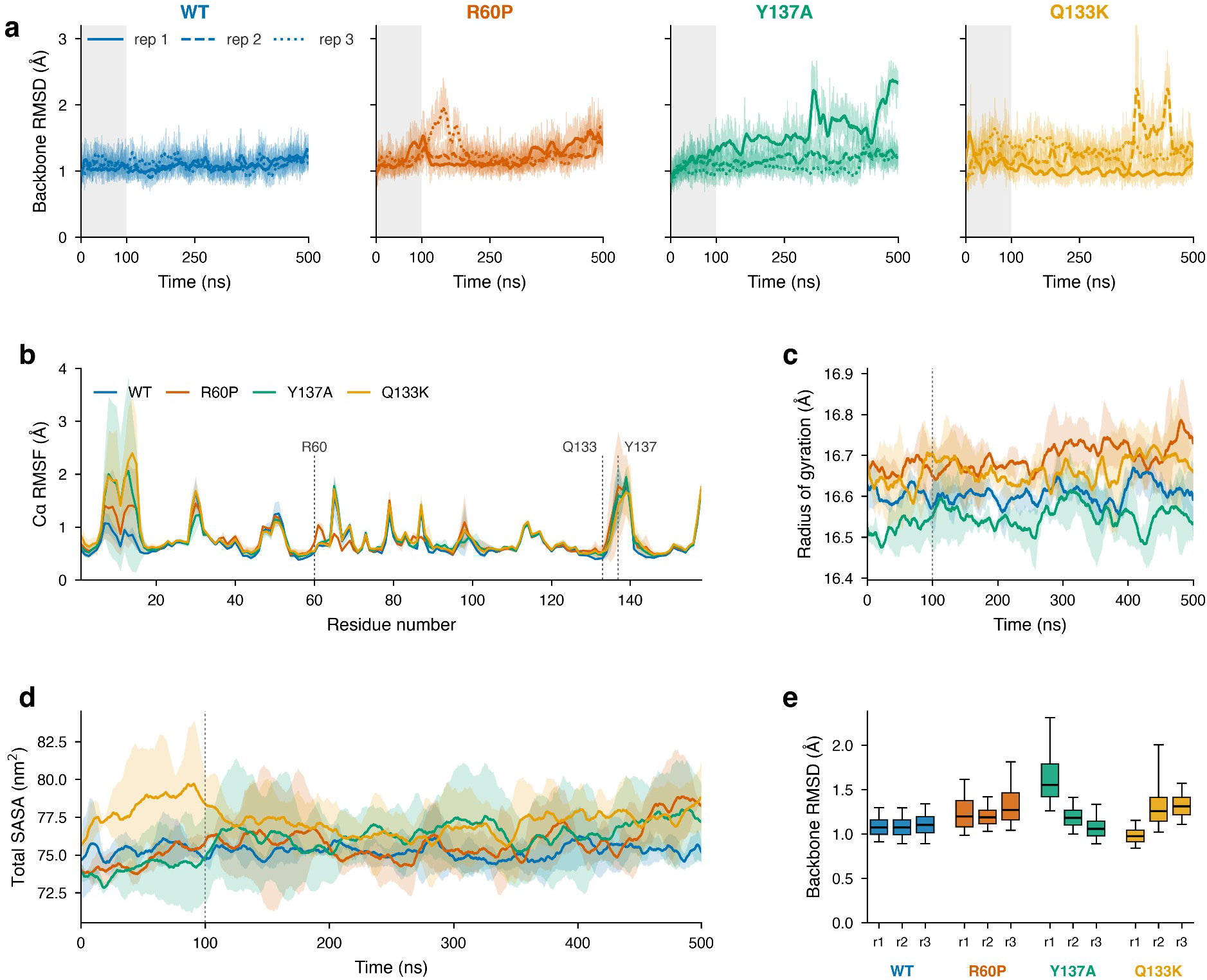
Global structural dynamics of wild-type FimH and the R60P, Y137A and Q133K variants (cpptraj pipeline; colours WT blue, R60P orange, Y137A green, Q133K gold throughout). (a) Backbone (N, Cα, C) RMSD over time per replicate (thin lines, raw; thick lines, 10 ns moving average; grey band, discarded first 100 ns). (b) Per-residue Cα RMSF (mean ± 1 SD between replicates); dotted verticals mark residues 60, 133 and 137. (c) Radius of gyration (mean ± 1 SD). (d) Total solvent-accessible surface area in nm² (mean ± 1 SD). (e) Distribution of backbone RMSD per replicate over 100-500 ns. n = 3 replicates per system.

**Figure 3.**
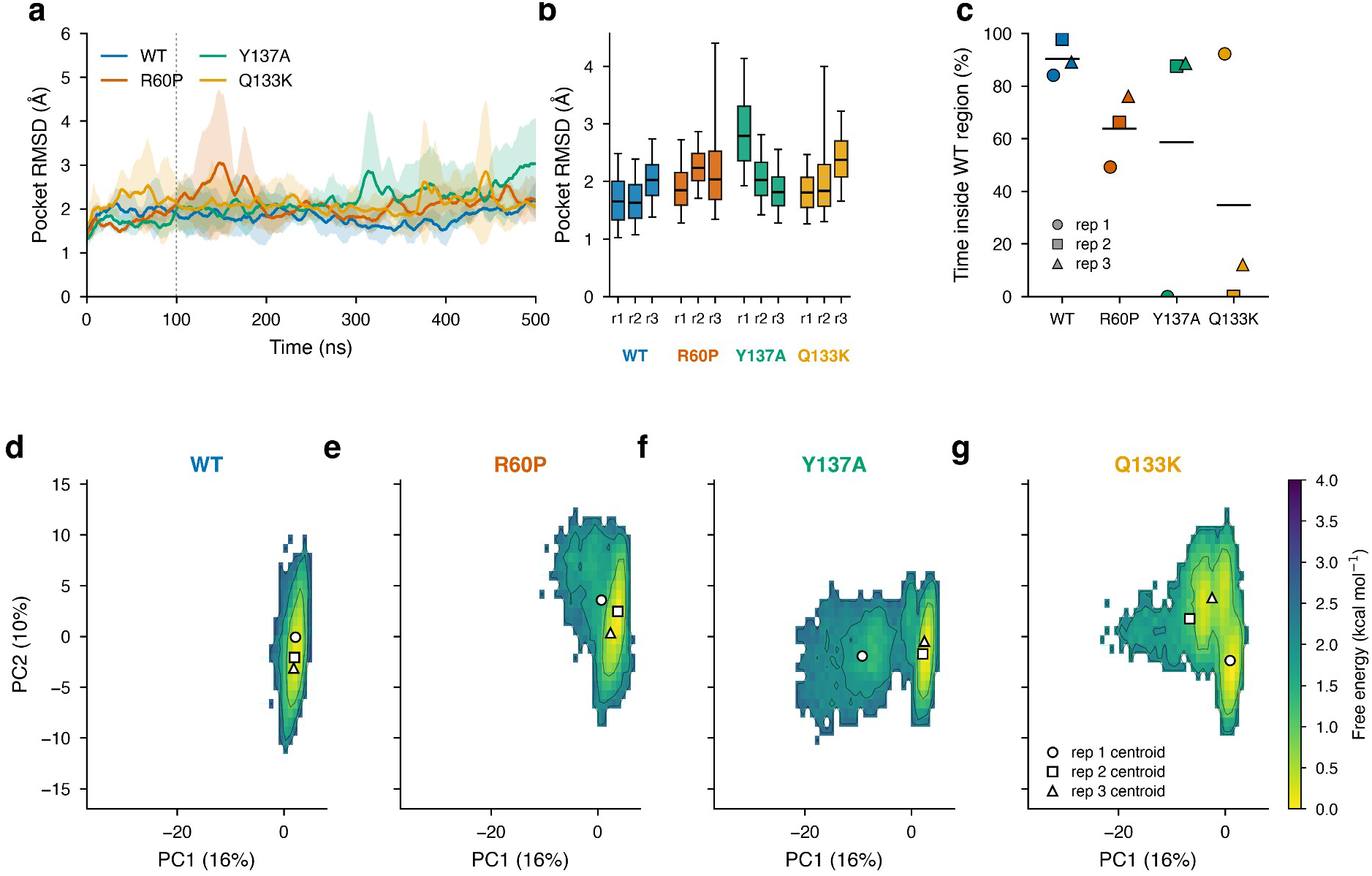
Binding-pocket dynamics and conformational landscape. (a) Pocket RMSD over time (thirteen binding-site residues; 10 ns moving average, mean ± SD of three replicates; dotted line, start of the analysis window). (b) Pocket RMSD per replicate over 100-500 ns. (c) Exploratory metric: percentage of each replicate’s frames inside the WT region of the PC1-PC2 landscape (cells with free energy ≤ 2 kcal/mol; horizontal bars, system mean). (d-g) Free-energy landscapes on PC1-PC2 (common covariance; ΔG = -k_B T ln P at 300 K) for WT, R60P, Y137A and Q133K, with the three replicate centroids. Colours WT blue, R60P orange, Y137A green, Q133K gold. n = 3 replicates per system.

**Figure 4.**
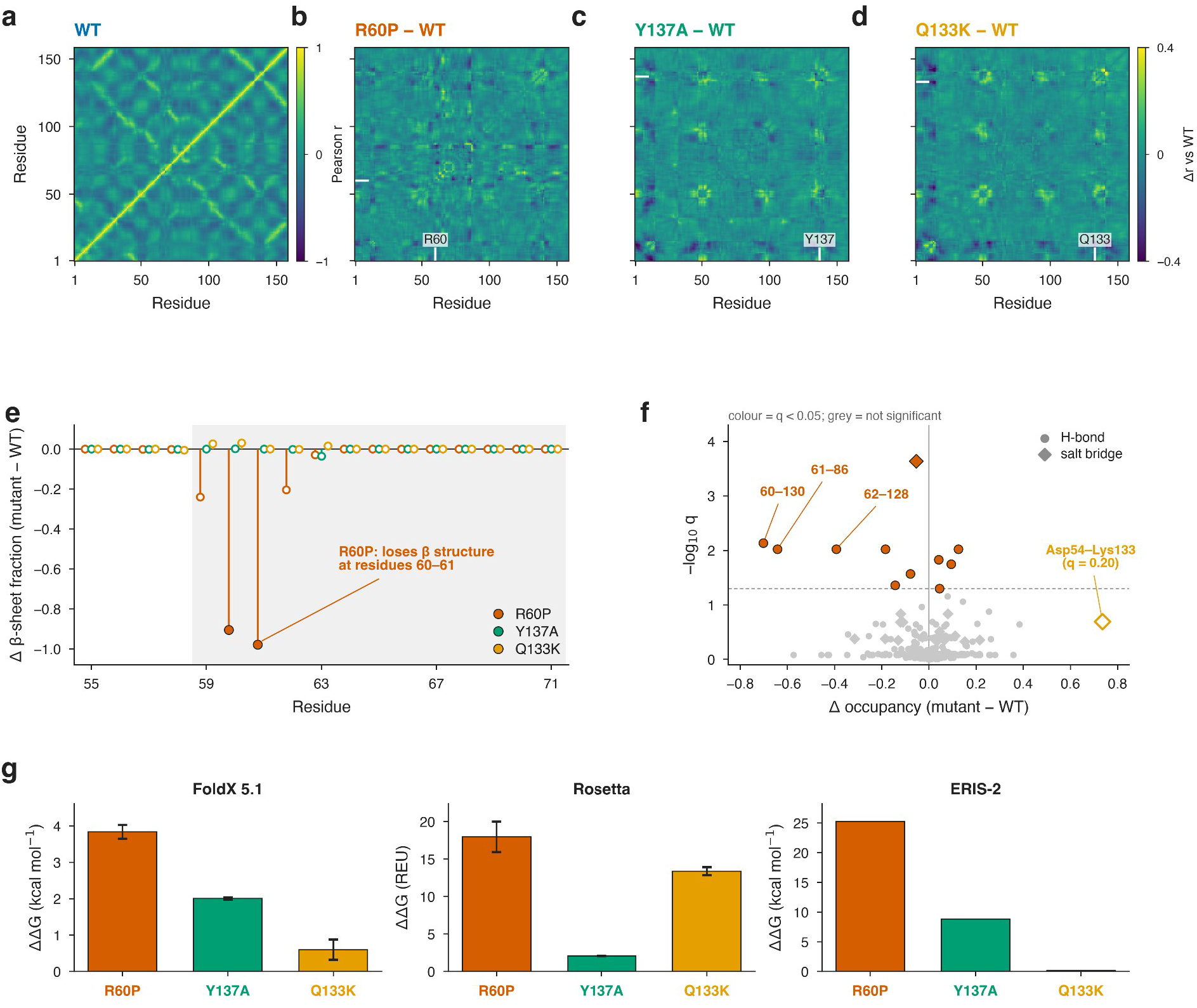
Correlated motions, local structure and predicted stability. (a) Dynamic cross-correlation map (DCCM) of WT (mean of three replicates, 100-500 ns; Pearson r, viridis colour scale). (b-d) Difference maps ΔDCCM = mutant − WT for R60P, Y137A and Q133K (viridis, fixed scale ±0.4); the mutated position is marked. (e) Change in beta-sheet fraction (mutant − WT) per residue over residues 55-71; filled symbols are significant (q < 0.05), open symbols are not; only R60P changes, at residues 60-61. (f) Volcano plot of the change in hydrogen-bond and salt-bridge occupancy versus −log10 q (exploratory); coloured points pass q < 0.05, grey points do not; the R60P hydrogen bonds 60-130, 61-86 and 62-128 and the exploratory Q133K Asp54-Lys133 salt bridge (q = 0.20) are labelled. (g) Predicted ΔΔG per method (FoldX, kcal/mol; Rosetta, REU; ERIS-2 classic, kcal/mol); note the different units (ERIS-2 classic input being confirmed; see Table 1). Colours WT blue, R60P orange, Y137A green, Q133K gold. n = 3 replicates per system for MD panels.

### 2.6 Statistics and quality control

The unit of analysis was the replicate (n = 3), because frames within a trajectory are autocorrelated; bars and bands denote the mean and ±1 SD between replicates. Differences from WT were expressed as a mean difference with a 95% confidence interval (Welch), and multiple comparisons were corrected by the Benjamini-Hochberg procedure (Benjamini & Hochberg, 1995). Because three replicates give low power, no hypothesis test was used to assert an effect; a difference was considered supported only when both the confidence interval excluded zero and the three mutant replicates fell outside the range of the three WT replicates. Effect thresholds (0.25 Angstrom for ΔRMSF, 0.20 for secondary-structure fraction, 0.25 for interaction occupancy, 0.30 for ΔDCCM), the 8 Angstrom shell and the salt-bridge definition (N-O < 4.0 Angstrom) were analytical decisions, not field standards. Hypotheses H1-H3 are the only confirmatory analyses and were fixed before the per-mutant trajectories were examined. The fraction of each replicate’s frames inside the WT region of the PC1-PC2 landscape (threshold 2 kcal/mol, cells of 0.9 units with a minimum of 10 frames; Figure 3c) and the interaction volcano plot (Figure 4f). Convergence followed best-practice recommendations (Grossfield et al., 2018): RMSIP between trajectory halves, cosine content of PC1-PC2, and the WT replicate-to-replicate control.

## 3. Results

### 3.1 Predicted stability

FoldX and ERIS-2 ranked the three variants in the same order, R60P > Y137A > Q133K, both placing R60P as the most destabilising and Q133K as effectively neutral (FoldX: 3.84 ± 0.19, 2.01 ± 0.03 and 0.60 ± 0.28 kcal/mol; ERIS-2 classic: 25.23, 8.82 and 0.21 kcal/mol). The Rosetta protocol also placed R60P highest (17.97 ± 2.03 REU) and Y137A low (2.07 ± 0.02 REU) but returned an anomalously high value for Q133K (13.39 ± 0.53 REU), discordant with both other predictors and with the MD results below; it is treated as a probable artefact of the custom protocol and is not interpreted as a real effect. The individual Rosetta values per relaxed structure are given in Table S1.

### 3.2 Global dynamics and convergence

All twelve trajectories stayed folded, with backbone RMSD fluctuating around 1.0-1.4 Angstrom after the discarded first 100 ns; WT was the most stable (1.10 ± 0.01 Angstrom, mean of replicates; Figure 2a,e). Two replicates departed from this baseline: Y137A replicate 1 drifted upward to about 2.5 Angstrom over the last 200 ns, and Q133K replicate 2 showed transient excursions near 3 Angstrom. The backbone-RMSD difference from WT excluded zero only for R60P (+0.155 Angstrom, 95% CI 0.029-0.282); for Y137A (+0.258 Angstrom) and Q133K (+0.124 Angstrom) the intervals were wide, driven by the divergent replicate, and included zero (Table 2). Per-residue RMSF (Figure 2b) was dominated by the mobile N-terminus and surface loops and was similar across systems, with no systematic shift at the mutated positions. The radius of gyration (Figure 2c) differed only slightly (R60P highest at 16.71 ± 0.05 Angstrom, Y137A lowest at 16.56 ± 0.05 Angstrom, WT 16.61 ± 0.01 Angstrom), and total SASA (Figure 2d, about 75-77 nm²) showed no supported difference.

**Table 2.** Quantitative MD summary per system over the 100-500 ns window, from the pipeline plotted in Figure 2. Mean ± SD are between the three replicates; the difference versus WT is shown with its 95% confidence interval (Welch). Bold marks the only difference whose interval excludes zero. SASA is given in nm² to match Figure 2d (1 nm² = 100 Å²).

| **Metric** | **System** | **Mean ± SD** | **Diff. vs WT** | **95% CI** |
|---|---|---|---|---|
| Backbone RMSD (Å) | WT | 1.099 ± 0.013 | — | — |
|  | R60P | 1.254 ± 0.054 | **+0.155** | 0.029 to 0.282 |
|  | Y137A | 1.356 ± 0.356 | +0.258 | -0.626 to 1.141 |
|  | Q133K | 1.222 ± 0.227 | +0.124 | -0.439 to 0.686 |
| Radius of gyration (Å) | WT | 16.612 ± 0.009 | — | — |
|  | R60P | 16.713 ± 0.050 | +0.101 | -0.016 to 0.219 |
|  | Y137A | 16.556 ± 0.046 | -0.056 | -0.165 to 0.053 |
|  | Q133K | 16.672 ± 0.046 | +0.060 | -0.049 to 0.168 |
| Total SASA (nm<sup>2</sup>) | WT | 75.2 ± 0.5 | — | — |
|  | R60P | 76.1 ± 1.4 | +0.8 | -2.3 to 4.0 |
|  | Y137A | 76.7 ± 2.5 | +1.5 | -4.4 to 7.4 |
|  | Q133K | 77.1 ± 1.7 | +1.9 | -2.2 to 5.9 |

### 3.3 Binding-pocket dynamics

Pocket RMSD over the thirteen binding-site residues rose from about 1.3 Angstrom to a plateau of 1.8-2.3 Angstrom in all systems (Figure 3a). Per replicate (Figure 3b), WT averaged 1.80 ± 0.21 Angstrom and the variants 2.11-2.26 Angstrom; the largest system mean, Y137A (2.26 ± 0.54 Angstrom), was produced by replicate 1 (2.88 Angstrom), whereas its other replicates (2.06 and 1.85 Angstrom) overlapped WT. R60P (2.15 ± 0.21 Angstrom) and Q133K (2.11 ± 0.29 Angstrom) were modestly higher than WT with overlapping distributions. Under the pre-set dual criterion no variant showed a supported increase in pocket RMSD, and the apparent Y137A effect is attributable to a single replicate. Results for the alternative 8 Angstrom shell definition are consistent and are given in the Supplementary Material.

### 3.4 Essential-dynamics landscape

A common-covariance PCA (PC1 and PC2 captured 16% and 10% of the variance) was used to compare the conformational space sampled by each system. As an exploratory metric, the fraction of each replicate’s frames falling inside the region populated by WT (free energy ≤ 2 kcal/mol in the pooled WT PC1-PC2 landscape; author-chosen threshold) was measured (Figure 3c). WT replicates stayed 84-98% inside this region. The variants were heterogeneous between replicates: R60P 49-76%; Y137A 0.0, 87.5 and 88.6%; and Q133K 92.2, 0.0 and 12.1% for replicates 1-3. The landscapes (Figure 3d-g) were a single basin for WT, a broader basin for R60P, two basins for Y137A — a second minimum near PC1 ≈ −10 populated only by replicate 1 — and a laterally extended basin for Q133K populated by replicates 2 and 3. In every case the departure from the WT landscape was produced by individual replicates rather than by a systematic, reproducible shift, and with three replicates per system it cannot be distinguished from replicate-to-replicate variability.

### 3.5 Correlated motions, secondary structure and contacts

Difference dynamic cross-correlation maps (ΔDCCM = mutant − WT; Figure 4b-d) showed sparse, mostly local changes; the significant disjoint pairs for R60P (for example 61-73, 61-86 and 83-87; Supplementary) lay near the mutated strand, whereas Y137A and Q133K changes were small and not robust to correction. The only reproduced secondary-structure change was in R60P, which lost beta-sheet content at residues 60 (Δ = −0.92) and 61 (Δ = −0.98) relative to WT (q < 0.05; Figure 4e); Y137A and Q133K were unchanged within replicate variability. Consistent with its strand disruption, R60P lost three local backbone hydrogen bonds — 60-130, 61-86 and 62-128 (q < 0.05; Figure 4f). For Q133K, the only notable contact change was conversion of the WT Gln133-Asp54 polar contact into a Lys133-Asp54 salt bridge present in about 74% of frames, but this was exploratory and did not survive correction (Δ = +0.74, q = 0.20; Figure 4f); no new interaction with Asp47 or Asp140 passed the threshold. No interaction change passed correction for Y137A.

## 4. Discussion

Across four systems simulated under identical conditions, a single lectin-domain substitution left the global fold intact and perturbed local structure to markedly different degrees. The clearest and most reproducible effect was the local structural perturbation at the proline substitution in R60P, accompanied by a consistently destabilising ΔΔG in both FoldX and classic ERIS-2. By contrast, the departures seen for Y137A and Q133K were produced by individual replicates — Y137A replicate 1 and Q133K replicates 2 and 3 in the free-energy landscape, and Y137A replicate 1 in the pocket RMSD — rather than by a systematic, reproducible effect. The strength of the present comparison lies in this controlled, replicate-based design across four systems and three independent stability predictors, which yields a coherent ranking of the variants and, for Q133K, a first dynamic characterisation.

Our support criterion is deliberately strict — a 95% confidence interval excluding zero together with non-overlapping replicate ranges — so “not established” below means not demonstrated at that threshold, not disproven. Against it, H1 is partially supported: R60P clearly altered local pocket structure (β-strand loss and lost hydrogen bonds), while the accompanying increase in flexibility was suggestive but did not reach the criterion. For H2, Thr158 did not enter the pocket on this timescale and no secondary-structure or interaction change reached threshold, so the hypothesis was not established here. For H3, the Lys133-Asp54 salt bridge is a genuine, reproducible rearrangement of the pre-existing Gln133-Asp54 contact, but it did not survive multiple-testing correction and no new Asp47/Asp140 contact appeared, so it remains a promising exploratory lead rather than an established result.

These observations sit within an established picture of FimH allostery, in which a soft central beta-sheet couples the interdomain region to the binding site (Interlandi & Thomas, 2016; Le Trong et al., 2010) and apparent affinity depends on binding-site populations hidden from the crystal (Languin-Cattoën et al., 2023). Residue 60 lies within the β-bulge/α-switch region (residues 59-71; Languin-Cattoën et al., 2023), so the local β-sheet loss we observe in R60P is a structural reading of where the substitution acts, not a demonstration of altered allostery. Our R60P result is consistent with reports that this substitution raises backbone fluctuations in the isolated lectin domain and serves as a minimal low-affinity model (Liu et al., 2021), although our replicate-based criterion is more conservative than single-trajectory comparisons and does not confirm a pocket-wide flexibility increase. For Y137A, Rabbani et al. (2017) reported from crystallography that Thr158 intrudes into the ligand-free pocket; we did not reproduce this on a 500 ns apo timescale, which may reflect the difference between a crystallographically trapped state and the dynamics accessible to a short simulation rather than a contradiction. For the previously uncharacterised Q133K, the conversion of a polar contact into a salt bridge is a plausible local mechanism but remains tentative.

The stability predictions ranked the variants consistently between FoldX and classic ERIS-2 (R60P > Y137A > Q133K) and identified R60P as the clearest destabiliser, matching the variant with the clearest MD perturbation; the Rosetta protocol agreed except for its anomalously high Q133K value, best read as a limitation of the custom protocol for a charge-changing substitution. Because the methods differ in units and input structure, we interpret only this rank order, and the ERIS-2 classic calculation will be repeated on the same input used for FoldX to confirm it. Importantly, folding-stability ΔΔG is not mannose affinity: a destabilisation of the apo monomer need not change binding or adhesion.

This study has limitations that define its scope. Three replicates of 500 ns provide a focused, hypothesis-driven comparison rather than a high-power screen, so some differences are suggestive without reaching our strict support criterion; the design is nonetheless sufficient to identify the reproducible R60P effect and to separate it from replicate-level variability. The work concerns the isolated apo lectin domain, without pilin, ligand or applied force, so it speaks to folding stability and local dynamics rather than to mannose affinity or adhesion. The R60P and Q133K substitutions also change the net charge, which cannot be fully separated from the structural change, and a single force field was used. Finally, the predictions are computational and await experimental validation. These constraints are typical of comparative molecular-dynamics studies and bound the interpretation without undermining the central, reproducible findings.

Several extensions would build directly on these results. Additional replicates or enhanced sampling would sharpen the distinction between reproducible effects and replicate-level variability, particularly for the suggestive flexibility changes. Simulating the complete lectin-plus-pilin construct under force would connect these local observations to the catch-bond, and simulating the variants in complex with mannose, with binding free-energy estimates, would test whether the predicted stability changes translate into affinity. Experimental validation by thermal-shift or calorimetric stability assays and adhesion measurements would anchor the computational picture, and confirming the stability predictions on a single, common input structure would further strengthen the cross-method agreement. Natural variants such as those of the ST131 lineage (Qin et al., 2022) provide a biologically grounded set of further mutations for the same framework.

## 5. Conclusions

Under identical, replicate-controlled conditions, the three FimH lectin-domain substitutions differed sharply in their consequences. Only R60P produced a clearly reproduced local effect at the mutated site and was also the variant most consistently flagged as destabilising by FoldX and classic ERIS-2, partially supporting H1; neither Y137A nor Q133K met its pre-specified criterion, leaving H2 and H3 unsupported. The departures of Y137A and Q133K from the wild-type conformational landscape were produced by individual replicates, not by a systematic shift. Predicted folding stability ranked the variants consistently between two methods but mapped only loosely onto the observed dynamics. Beyond these specific findings, the study offers a reproducible framework — with pre-specified thresholds, an inter-replicate control, and a clear separation of confirmatory from exploratory analyses — for interpreting single FimH mutations.

## Declarations

### Data and code availability

All supporting data and code are provided in a Supplementary ZIP archive: starting structures (PDB 4XO8 and mutant models), system topologies and coordinates, molecular dynamics input files with random seeds, raw per-replicate time series and energy outputs (FoldX, Rosetta, ERIS-2), source tables, analysis scripts and figure-generation code.

### Author contributions

J.A.N.-P.: Conceptualization, Methodology, Software, Formal analysis, Investigation, Data curation, Visualization, Writing – original draft. A.P.-T.: Conceptualization, Validation, Resources, Writing – review & editing. C.V.-D.: Conceptualization, Validation, Resources, Writing – review & editing. A.A.-A.: Conceptualization, Methodology, Validation, Supervision, Project administration, Writing – review & editing. All authors read and approved the final manuscript.

### Use of AI tools

The authors certify that no AI-based tools or applications were used throughout this research effort starting from its inception to data analysis. Only the work of the authors is reflected in this study’s methodology and results. Although the authors did use an AI-based linguistic enhancement application to improve syntax, readability and overall cohesiveness of the document after the initial draft of the manuscript was completed, the authors reviewed and revised the recommendations provided by the software and assume full accountabilities for all aspects of the final version of the manuscript and its academic integrity

## Supporting information

Metadata

## References

Åqvist, J., Wennerström, P., Nervall, M., Bjelic, S., & Brandsdal, B. O. (2004). Molecular dynamics simulations of water and biomolecules with a Monte Carlo constant pressure algorithm. Chemical Physics Letters, 384(4-6), 288–294. 10.1016/j.cplett.2003.12.039

Benjamini, Y., & Hochberg, Y. (1995). Controlling the false discovery rate: A practical and powerful approach to multiple testing. Journal of the Royal Statistical Society: Series B, 57(1), 289–300. 10.1111/j.2517-6161.1995.tb02031.x

Berendsen, H. J. C., Postma, J. P. M., van Gunsteren, W. F., DiNola, A., & Haak, J. R. (1984). Molecular dynamics with coupling to an external bath. The Journal of Chemical Physics, 81(8), 3684–3690. 10.1063/1.448118

Case, D. A., Aktulga, H. M., Belfon, K., Cerutti, D. S., Cisneros, G. A., Cruzeiro, V. W. D., … Merz, K. M. (2023). AmberTools. Journal of Chemical Information and Modeling, 63(20), 6183–6191. 10.1021/acs.jcim.3c01153

Cock, P. J. A., Antao, T., Chang, J. T., Chapman, B. A., Cox, C. J., Dalke, A., … de Hoon, M. J. L. (2009). Biopython: Freely available Python tools for computational molecular biology and bioinformatics. Bioinformatics, 25(11), 1422–1423. 10.1093/bioinformatics/btp163

Darden, T., York, D., & Pedersen, L. (1993). Particle mesh Ewald: An N·log(N) method for Ewald sums in large systems. The Journal of Chemical Physics, 98(12), 10089–10092. 10.1063/1.464397

Eastman, P., Swails, J., Chodera, J. D., McGibbon, R. T., Zhao, Y., Beauchamp, K. A., … Pande, V. S. (2017). OpenMM 7: Rapid development of high performance algorithms for molecular dynamics. PLOS Computational Biology, 13(7), e1005659. 10.1371/journal.pcbi.1005659

Grossfield, A., Patrone, P. N., Roe, D. R., Schultz, A. J., Siderius, D. W., & Zuckerman, D. M. (2018). Best practices for quantification of uncertainty and sampling quality in molecular simulations. Living Journal of Computational Molecular Science, 1(1), 5067. 10.33011/livecoms.1.1.5067

Hatton, N. E., Baumann, C. G., & Fascione, M. A. (2021). Developments in mannose-based treatments for uropathogenic Escherichia coli-induced urinary tract infections. ChemBioChem, 22(4), 613–629. 10.1002/cbic.202000406

Hopkins, C. W., Le Grand, S., Walker, R. C., & Roitberg, A. E. (2015). Long-time-step molecular dynamics through hydrogen mass repartitioning. Journal of Chemical Theory and Computation, 11(4), 1864–1874. 10.1021/ct5010406

Interlandi, G., & Thomas, W. E. (2016). Mechanism of allosteric propagation across a β-sheet structure investigated by molecular dynamics simulations. Proteins, 84(7), 990–1008. 10.1002/prot.25050

Jorgensen, W. L., Chandrasekhar, J., Madura, J. D., Impey, R. W., & Klein, M. L. (1983). Comparison of simple potential functions for simulating liquid water. The Journal of Chemical Physics, 79(2), 926–935. 10.1063/1.445869

Joung, I. S., & Cheatham, T. E., III. (2008). Determination of alkali and halide monovalent ion parameters for use in explicitly solvated biomolecular simulations. The Journal of Physical Chemistry B, 112(30), 9020–9041. 10.1021/jp8001614

Kabsch, W., & Sander, C. (1983). Dictionary of protein secondary structure: Pattern recognition of hydrogen-bonded and geometrical features. Biopolymers, 22(12), 2577–2637. 10.1002/bip.360221211

Languin-Cattoën, O., Sterpone, F., & Stirnemann, G. (2023). Binding site plasticity regulation of the FimH catch-bond mechanism. Biophysical Journal, 122(13), 2744–2756. 10.1016/j.bpj.2023.05.029

Le Trong, I., Aprikian, P., Kidd, B. A., Forero-Shelton, M., Tchesnokova, V., Rajagopal, P., … Thomas, W. E. (2010). Structural basis for mechanical force regulation of the adhesin FimH via finger trap-like β sheet twisting. Cell, 141(4), 645–655. 10.1016/j.cell.2010.03.038

Li, P., Song, L. F., & Merz, K. M., Jr. (2015). Systematic parameterization of monovalent ions employing the nonbonded model. Journal of Chemical Theory and Computation, 11(4), 1645–1657. 10.1021/ct500918t

Li, X., Fan, H., Zi, H., Hu, H., Li, B., Huang, J., … Zeng, X. (2022). Global and regional burden of bacterial antimicrobial resistance in urinary tract infections in 2019. Journal of Clinical Medicine, 11(10), 2817. 10.3390/jcm11102817

Liu, J., Amaral, L. A. N., & Keten, S. (2021). Conformational stability of the bacterial adhesin, FimH, with an inactivating mutation. Proteins, 89(3), 276–288. 10.1002/prot.26013

Maier, J. A., Martinez, C., Kasavajhala, K., Wickstrom, L., Hauser, K. E., & Simmerling, C. (2015). ff14SB: Improving the accuracy of protein side chain and backbone parameters from ff99SB. Journal of Chemical Theory and Computation, 11(8), 3696–3713. 10.1021/acs.jctc.5b00255

Mancuso, G., Midiri, A., Gerace, E., Marra, M., Zummo, S., & Biondo, C. (2023). Urinary tract infections: The current scenario and future prospects. Pathogens, 12(4), 623. 10.3390/pathogens12040623

Mir, S. A., Nayak, B., Aljarba, N. H., Kumarasamy, V., Subramaniyan, V., & Dhara, B. (2024). Exploring KRas protein dynamics: An integrated molecular dynamics analysis of KRas wild and mutant variants. ACS Omega, 9(28), 30665–30674. 10.1021/acsomega.4c02671

Park, H., Bradley, P., Greisen, P., Jr., Liu, Y., Mulligan, V. K., Kim, D. E., … DiMaio, F. (2016). Simultaneous optimization of biomolecular energy functions on features from small molecules and macromolecules. Journal of Chemical Theory and Computation, 12(12), 6201–6212. 10.1021/acs.jctc.6b00819

Pettersen, E. F., Goddard, T. D., Huang, C. C., Meng, E. C., Couch, G. S., Croll, T. I., Morris, J. H., & Ferrin, T. E. (2021). UCSF ChimeraX: Structure visualization for researchers, educators, and developers. Protein Science, 30(1), 70–82. 10.1002/pro.3943

Qin, J., Wilson, K. A., Sarkar, S., Heras, B., O’Mara, M. L., & Totsika, M. (2022). Conserved FimH mutations in the global Escherichia coli ST131 multi-drug resistant lineage weaken interdomain interactions and alter adhesin function. Computational and Structural Biotechnology Journal, 20, 4532–4541. 10.1016/j.csbj.2022.08.040

Rabbani, S., Krammer, E.-M., Roos, G., Zalewski, A., Preston, R., Eid, S., … Bouckaert, J. (2017). Mutation of Tyr137 of the universal Escherichia coli fimbrial adhesin FimH relaxes the tyrosine gate prior to mannose binding. IUCrJ, 4(1), 7–23. 10.1107/S2052252516016675

Redler, R. L., Das, J., Diaz, J. R., & Dokholyan, N. V. (2016). Protein destabilization as a common factor in diverse inherited disorders. Journal of Molecular Evolution, 82(1), 11–16. 10.1007/s00239-015-9717-5

Roe, D. R., & Cheatham, T. E., III. (2013). PTRAJ and CPPTRAJ: Software for processing and analysis of molecular dynamics trajectory data. Journal of Chemical Theory and Computation, 9(7), 3084–3095. 10.1021/ct400341p

Ryckaert, J.-P., Ciccotti, G., & Berendsen, H. J. C. (1977). Numerical integration of the Cartesian equations of motion of a system with constraints: Molecular dynamics of n-alkanes. Journal of Computational Physics, 23(3), 327–341. 10.1016/0021-9991(77)90098-5

Sapozhnikov, Y., Patel, J. S., Ytreberg, F. M., & Miller, C. R. (2023). Statistical modeling to quantify the uncertainty of FoldX-predicted protein folding and binding stability. BMC Bioinformatics, 24(1), 426. 10.1186/s12859-023-05537-0

Sarshar, M., Behzadi, P., Ambrosi, C., Zagaglia, C., Palamara, A. T., & Scribano, D. (2020). FimH and anti-adhesive therapeutics: A disarming strategy against uropathogens. Antibiotics, 9(7), 397. 10.3390/antibiotics9070397

Sauer, M. M., Jakob, R. P., Eras, J., Baday, S., Eriş, D., Navarra, G., … Glockshuber, R. (2016). Catch-bond mechanism of the bacterial adhesin FimH. Nature Communications, 7, 10738. 10.1038/ncomms10738

Schymkowitz, J., Borg, J., Stricher, F., Nys, R., Rousseau, F., & Serrano, L. (2005). The FoldX web server: An online force field. Nucleic Acids Research, 33(Web Server issue), W382–W388. 10.1093/nar/gki387

Thiltgen, G., & Goldstein, R. A. (2012). Assessing predictors of changes in protein stability upon mutation using self-consistency. PLOS ONE, 7(10), e46084. 10.1371/journal.pone.0046084

Thomas, W. E., Trintchina, E., Forero, M., Vogel, V., & Sokurenko, E. V. (2002). Bacterial adhesion to target cells enhanced by shear force. Cell, 109(7), 913–923. 10.1016/S0092-8674(02)00796-1

Whelan, S., Lucey, B., & Finn, K. (2023). Uropathogenic Escherichia coli (UPEC)-associated urinary tract infections: The molecular basis for challenges to effective treatment. Microorganisms, 11(9), 2169. 10.3390/microorganisms11092169

Williams, B., II, Convertino, M., Das, J., & Dokholyan, N. V. (2017). Molecular mechanisms of the R61T mutation in apolipoprotein E4: A dynamic rescue. Biophysical Journal, 113(10), 2192–2198. 10.1016/j.bpj.2017.08.026

Zhang, D. Y., Wang, J., & Dokholyan, N. V. (2021). Prefusion spike protein stabilization through computational mutagenesis. Proteins, 89(4), 399–408. 10.1002/prot.26025

