## Supplementary material for "Point mutations in the FimH lectin domain: predicted folding stability versus conformational dynamics for R60P, Q133K and Y137A": Metadata: Figura1_panel.pdf

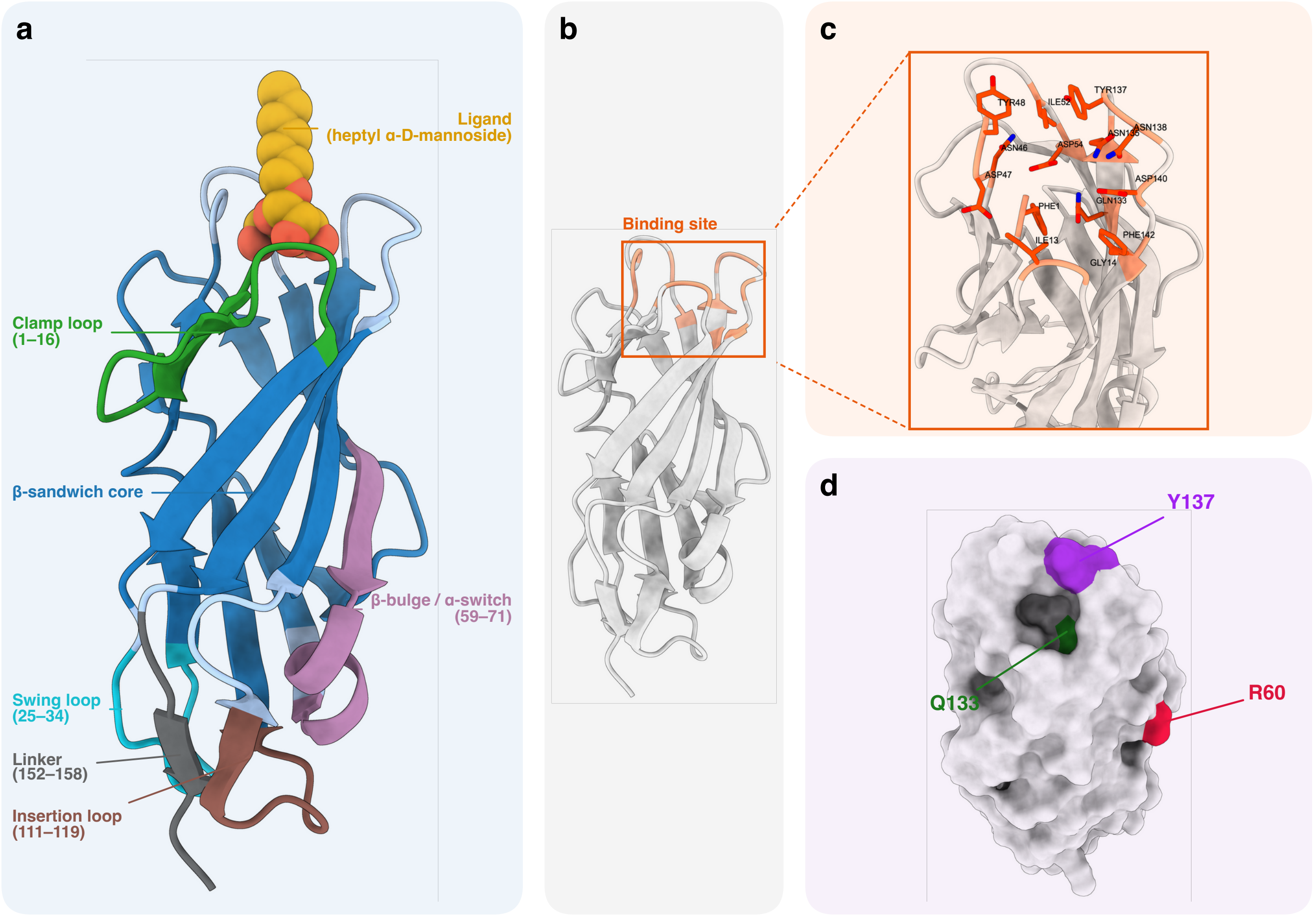
a
Ligand
(heptyl α-D-mannoside)
Clamp loop
(1–16)
β-sandwich core
β-bulge / α-switch
(59–71)
Swing loop
(25–34)
Linker
(152–158)
Insertion loop
(111–119)
b
Binding site
c
TYR48
ILE52
TYR137
ASN138
ASN135
ASN46
ASP54
ASP47
ASP140
GLN133
PHE1
PHE142
ILE13
GLY14
d
Y137
Q133
R60
