## Supplementary material for "Point mutations in the FimH lectin domain: predicted folding stability versus conformational dynamics for R60P, Q133K and Y137A": Metadata: Figure2_panel.pdf

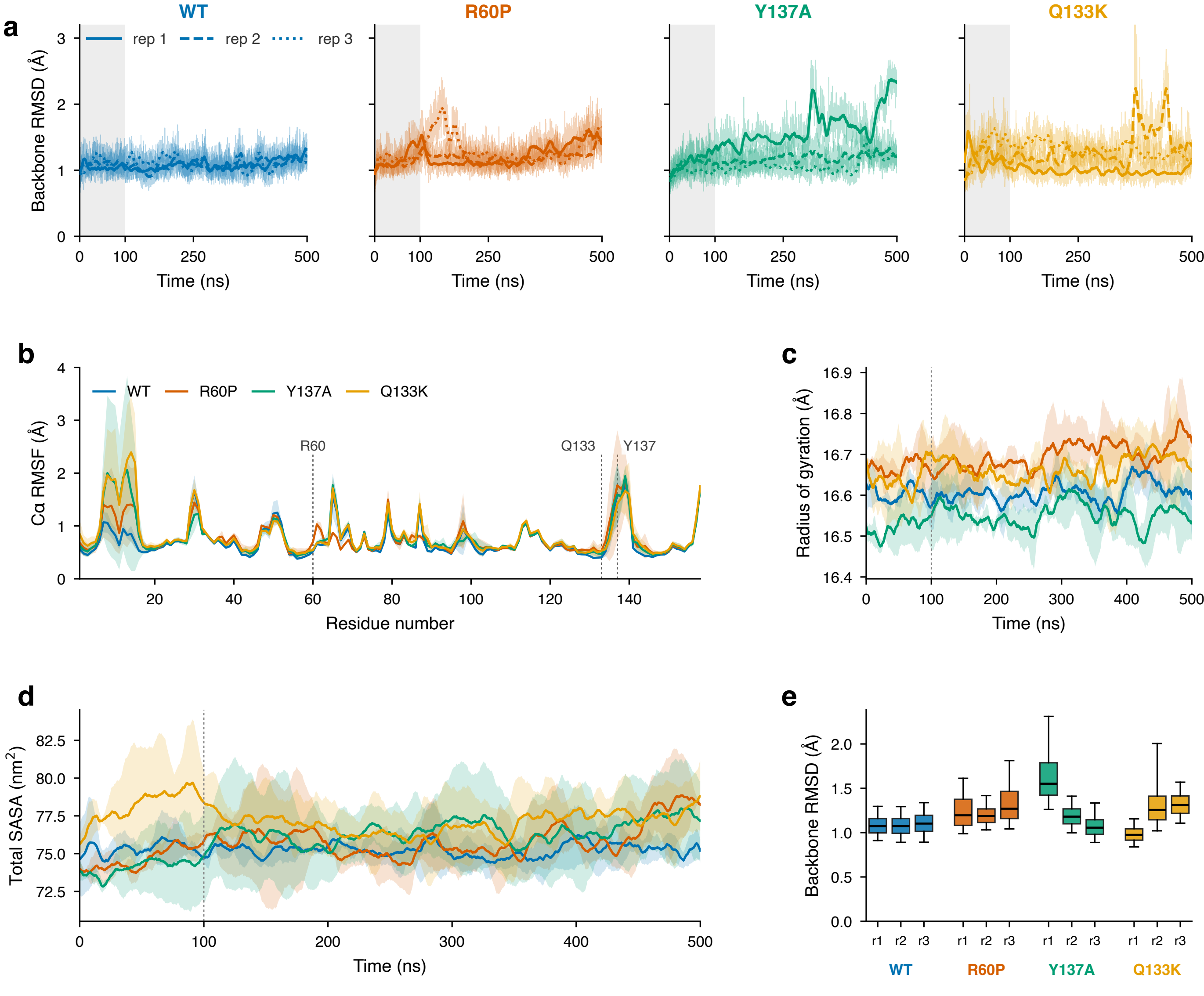

a
WT
R60P
Y137A
Q133K
rep 1
rep 2
rep 3
Backbone RMSD (Å)
Time (ns)
b
Cα RMSF (Å)
Residue number
R60
Q133
Y137
c
Radius of gyration (Å)
Time (ns)
d
Total SASA (nm$^2$)
Time (ns)
e
Backbone RMSD (Å)
r1
r2
r3
