## Supplementary material for "Point mutations in the FimH lectin domain: predicted folding stability versus conformational dynamics for R60P, Q133K and Y137A": Metadata: Figure3_panel.pdf

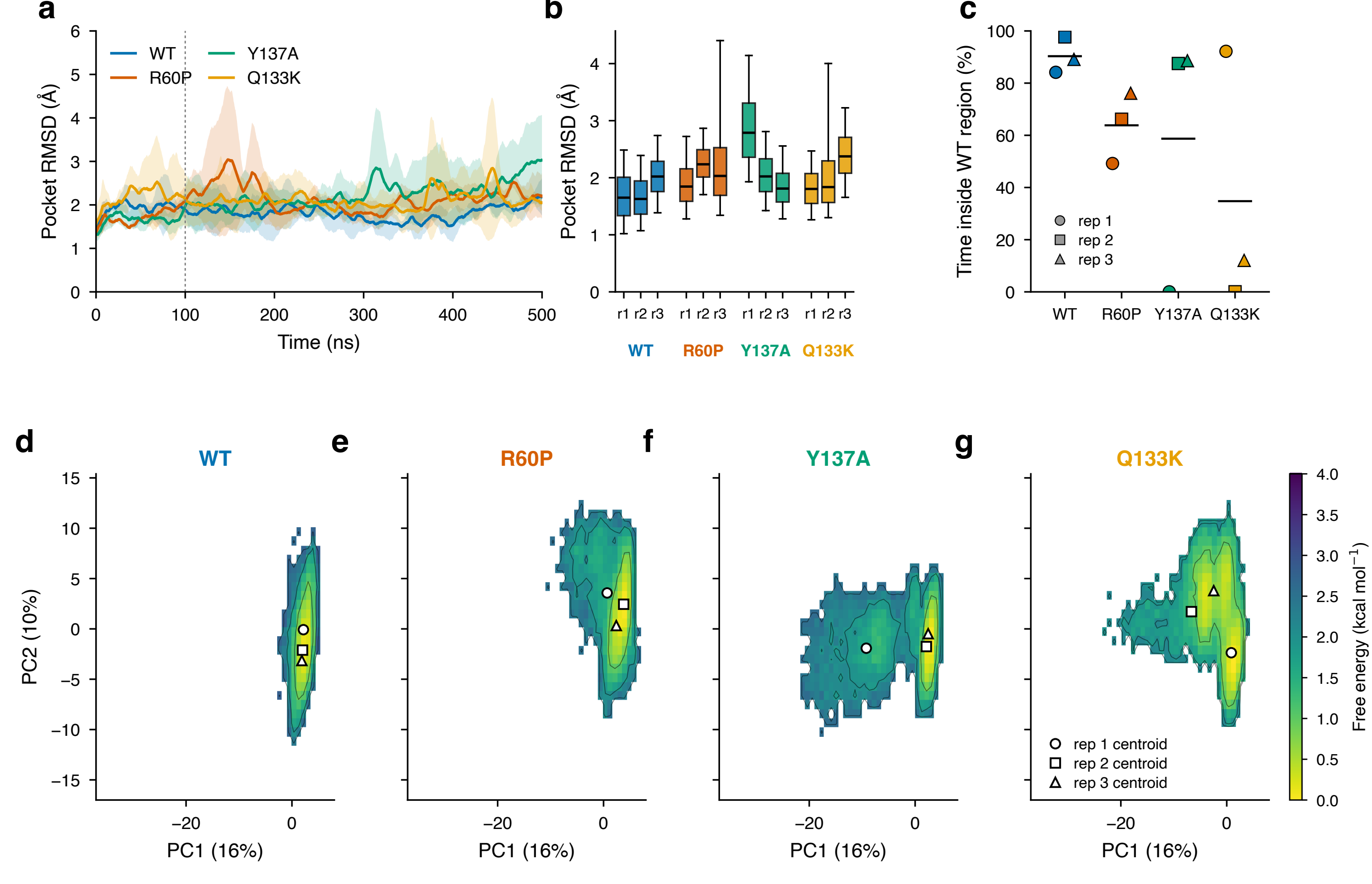
a
b
c
d
e
f
g
WT
R60P
Y137A
Q133K
Pocket RMSD (Å)
Time (ns)
r1 r2 r3
Time inside WT region (%)
rep 1
rep 2
rep 3
PC2 (10%)
PC1 (16%)
rep 1 centroid
rep 2 centroid
rep 3 centroid
Free energy (kcal mol$^{-1}$)
