## Supplementary material for "Point mutations in the FimH lectin domain: predicted folding stability versus conformational dynamics for R60P, Q133K and Y137A": Metadata: Figure5_panel.pdf

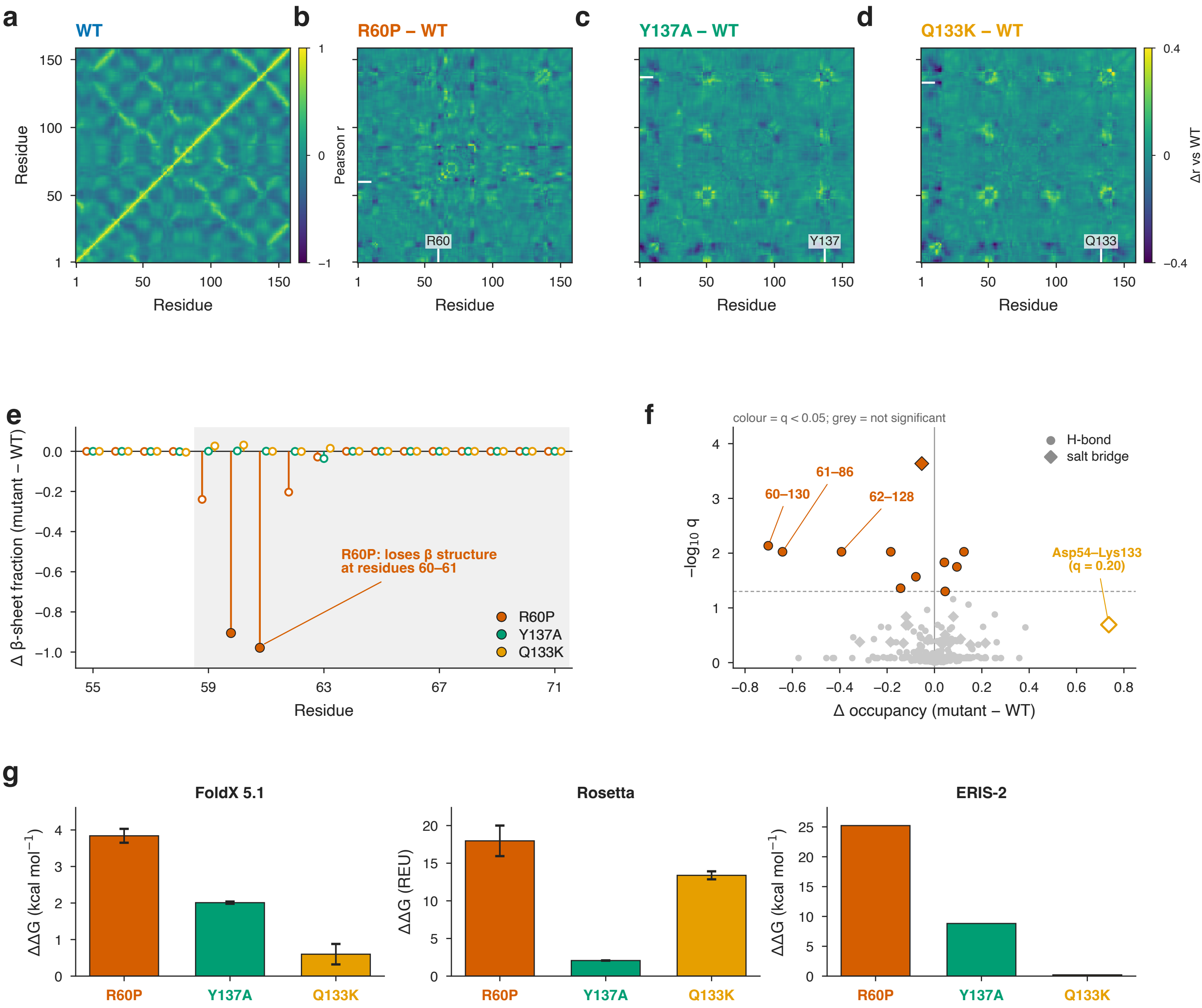

a
WT
Residue
Pearson r
b
R60P – WT
R60
c
Y137A – WT
Y137
d
Q133K – WT
Q133
Δr vs WT
Residue
e
Δ β-sheet fraction (mutant – WT)
R60P: loses β structure at residues 60–61
R60P
Y137A
Q133K
Residue
f
colour = q < 0.05; grey = not significant
−log10 q
H-bond
salt bridge
61–86
60–130
62–128
Asp54–Lys133 (q = 0.20)
Δ occupancy (mutant – WT)
g
FoldX 5.1
ΔΔG (kcal mol−1)
Rosetta
ΔΔG (REU)
ERIS-2
R60P
Y137A
Q133K
