## Supplementary material for "Point mutations in the FimH lectin domain: predicted folding stability versus conformational dynamics for R60P, Q133K and Y137A": Metadata: Suplementarios_FimH.docx

**Figure S1 (former per-residue ΔRMSF).** Per-residue difference in Cα RMSF (mutant − WT) with 95% confidence intervals; no residue met the pre-set criterion (q_BH < 0.05 and |ΔRMSF| ≥ 0.25 Angstrom) in any variant. This analysis is retained as supplementary because per-residue RMSF is shown in main Figure 2b. Source: dRMSF_<mutant>_vs_WT.csv.

**8 Angstrom shell pocket RMSD.** Using the alternative shell definition (all residues within 8 Angstrom of each mutated site), local-fit RMSD differences from WT were +0.216 (R60P), +0.161 (Y137A) and +0.316 Angstrom (Q133K); all 95% confidence intervals included zero, consistent with the 13-residue pocket analysis in Figure 3a-b.

**ERIS-2 neural-mode second opinion.** A neural-mode ERIS-2 run returned R60P 0.01, Y137A 0.15 and Q133K 0.46 kcal/mol (all classified neutral), differing from the classic-mode values in Table 1. The input structures of both ERIS-2 runs are being confirmed, and only the classic-mode rank order is used in the main text.

**Significant ΔDCCM pairs and full interaction table.** Disjoint significant ΔDCCM pairs (|Δ| ≥ 0.30) per mutant and the complete per-pair hydrogen-bond and salt-bridge table (occupancy, Δ, q) are provided as data files (dDCCM_<mutant>_pairs_absdelta_ge_0.3.csv; interactions_<mutant>_vs_WT.csv).

**R60P proline-ring reconstruction.** In R60P, PDBFixer left the Pro60 ring open (N-Cδ = 4.53 Angstrom); the Cγ and Cδ atoms were rebuilt by internal coordinates (N-Cδ = 1.57 Angstrom after correction) and the geometry held throughout the trajectories (N-Cδ = 1.44-1.50 Angstrom).

**Equilibration protocol.** The nine-stage restrained equilibration summarised in Methods 2.3 is detailed stage by stage in Table S3.

**Trajectory quality control.** Each 50 ns segment yielded 5000 frames at Δt = 10 ps with no gaps and no NaN values; the temperature stayed within 292.6-307.8 K without drift, the density within 1.002-1.028 g/cm³, and no trajectory showed denaturation.

**Software versions.** Amber 26 and AmberTools 24.8 (pmemd.cuda, tleap, cpptraj 6.24), PDBFixer 1.12, OpenMM 8.6.1, Biopython 1.88 and Python 3.11.18; production ran on an NVIDIA RTX 5070 Ti at about 247 ns/day per run.

**R60P 132-141 salt bridge.** This salt bridge changed significantly (q ≈ 0.0002) but by a negligible amount (Δ occupancy -0.05) and is of no structural consequence; it is noted here for completeness.

**Table S1.** Individual Rosetta ΔΔG values (REU) per independently relaxed structure (custom Cartesian-ddG-type protocol). Means ± SD are reproduced in Table 1.

| Mutation | relax 0 | relax 1 | relax 2 | Mean ± SD |
| --- | --- | --- | --- | --- |
| R60P | 16.89 | 16.71 | 20.32 | 17.97 ± 2.03 |
| Q133K | 13.35 | 12.89 | 13.94 | 13.39 ± 0.53 |
| Y137A | 2.09 | 2.06 | 2.06 | 2.07 ± 0.02 |

**Table S2 (System composition).** Composition of the four simulated systems. Net charge is for the protein before adding ions; the initial box edge is the value at the start of production.

| System | Net charge | Atoms | Waters | Na+ | Cl- | [salt] (mM) | Box (Å) |
| --- | --- | --- | --- | --- | --- | --- | --- |
| WT | -1 | 30 254 | 9281 | 26 | 25 | 149 | 73.39 |
| R60P | -2 | 30 308 | 9302 | 27 | 25 | 149 | 73.37 |
| Y137A | -1 | 30 210 | 9270 | 26 | 25 | 150 | 73.20 |
| Q133K | 0 | 30 189 | 9258 | 25 | 25 | 150 | 73.33 |

**Table S3 (Equilibration protocol).** Nine-stage restrained equilibration. Common settings: 2 fs time step, 8.0 Angstrom cutoff, SHAKE on bonds to hydrogen, and position restraints on the heavy atoms of residues 1-158 referenced to the initial coordinates; stages 5-9 used a Berendsen thermostat (τ_T = 0.5 ps) and, for NPT, barostat (τ_P = 1.0 ps).

| Stage | Ensemble | Length | Restraint (kcal/mol/Å²) |
| --- | --- | --- | --- |
| 1 | Minimisation | 5000 cycles (2500 SD + 2500 CG) | 5.0 |
| 2 | Minimisation | 5000 cycles | 2.0 |
| 3 | Minimisation | 5000 cycles | 0.1 |
| 4 | Minimisation | 5000 cycles | 0 |
| 5 | NVT, 300 K | 50 ps | 5.0 |
| 6 | NPT, 1 atm | 100 ps | 1.0 |
| 7 | NPT | 100 ps | 0.5 |
| 8 | NPT | 100 ps | 0.5 on N, Cα, C, O, Cβ |
| 9 | NPT | 100 ps | none |
