## Supplementary figures and images for "Point mutations in the FimH lectin domain: predicted folding stability versus conformational dynamics for R60P, Q133K and Y137A"

### Figura1_panel.png

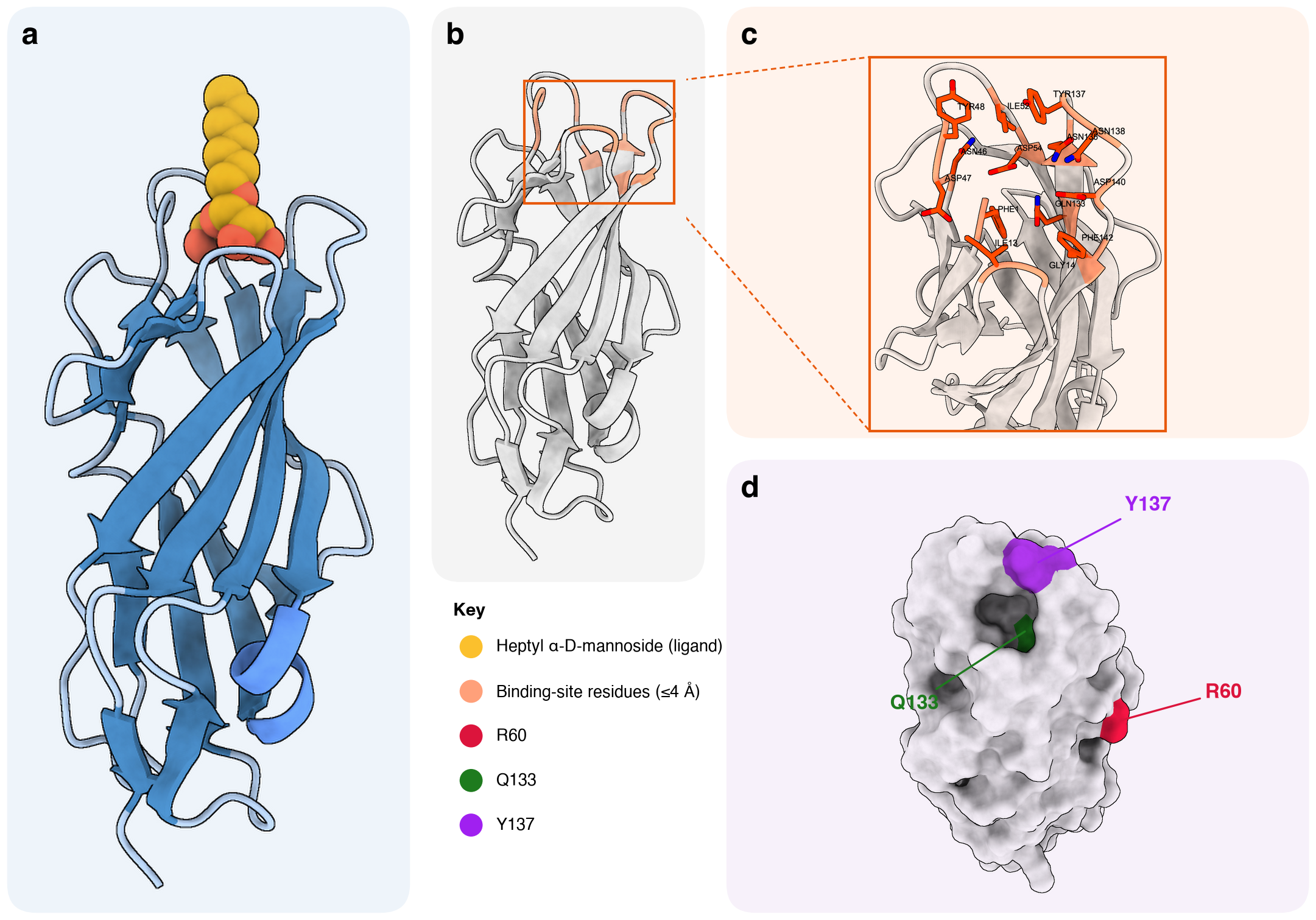

### Figura1_panel_600dpi.jpg

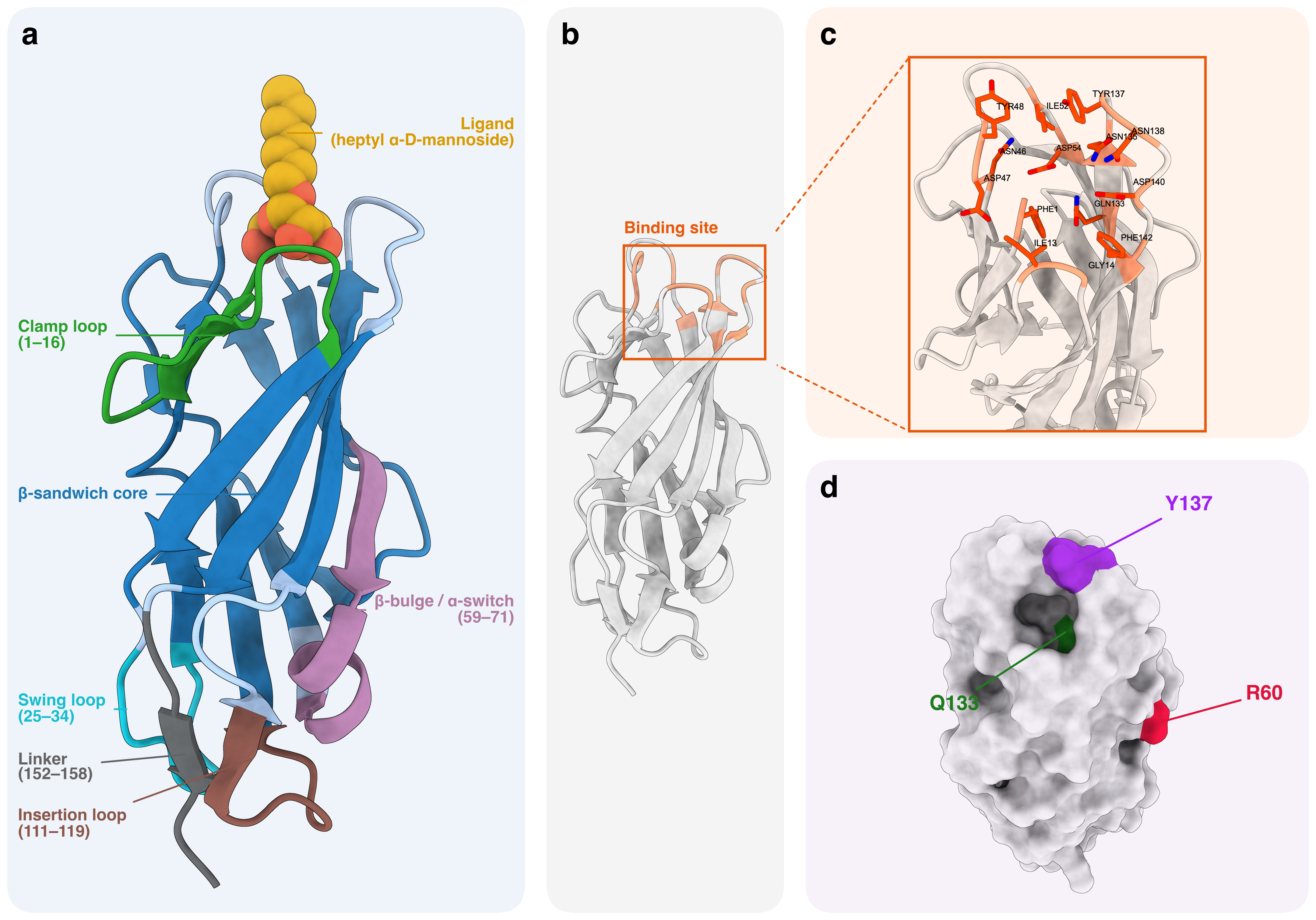

### Figura1_panel_600dpi.png

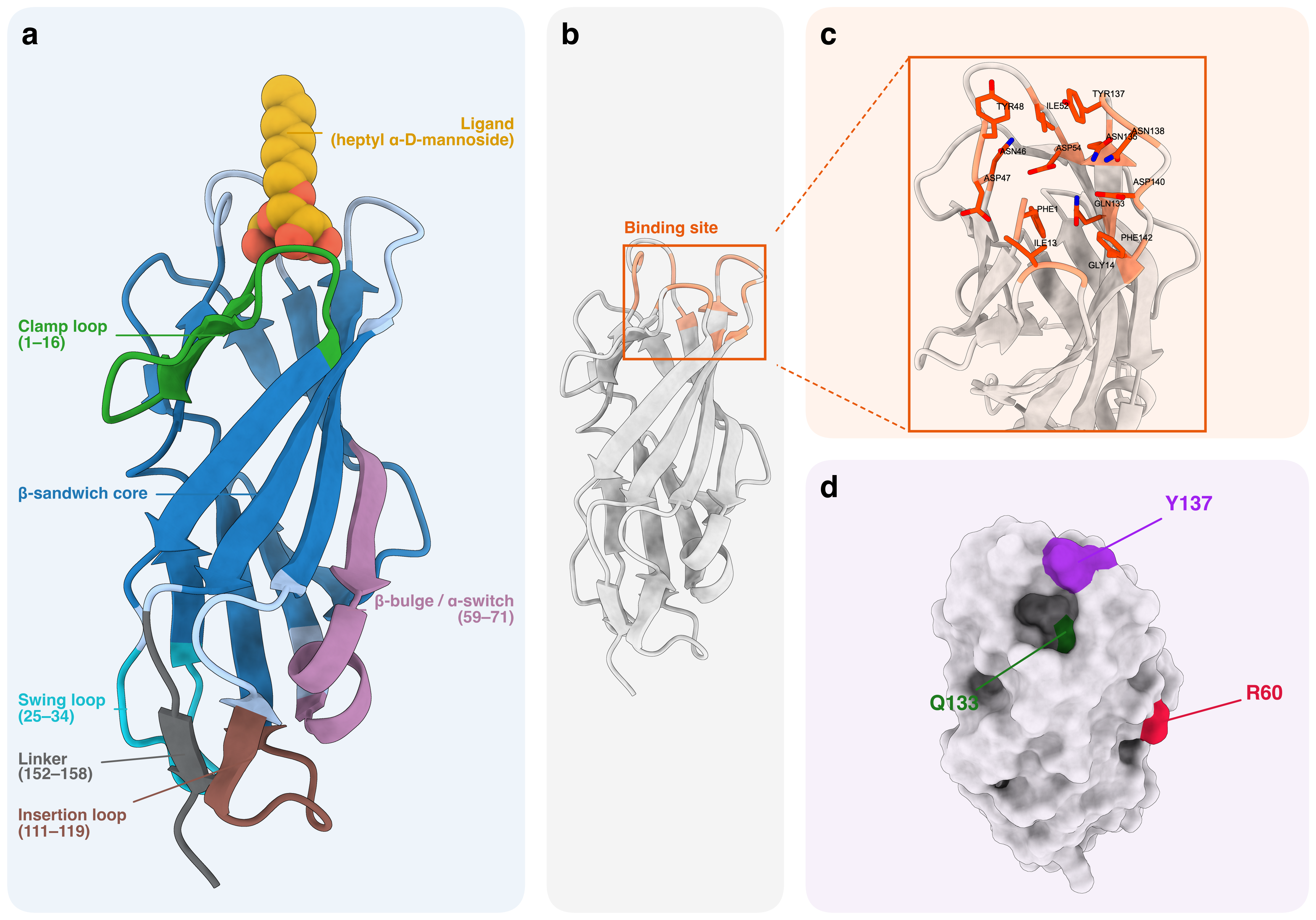

### Figura1_panel_1200dpi.tiff

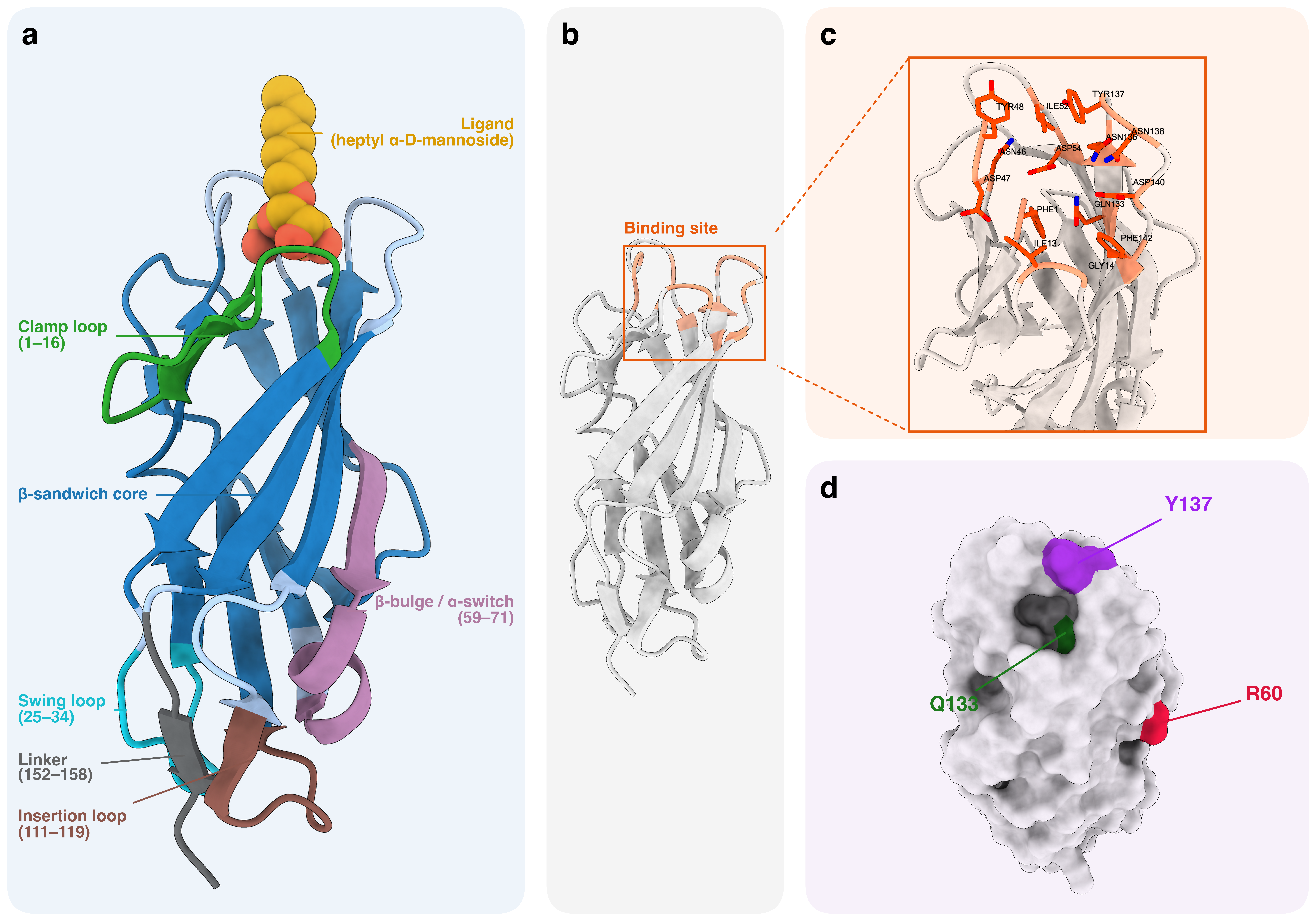

### Figure2_dinamica_global.jpg

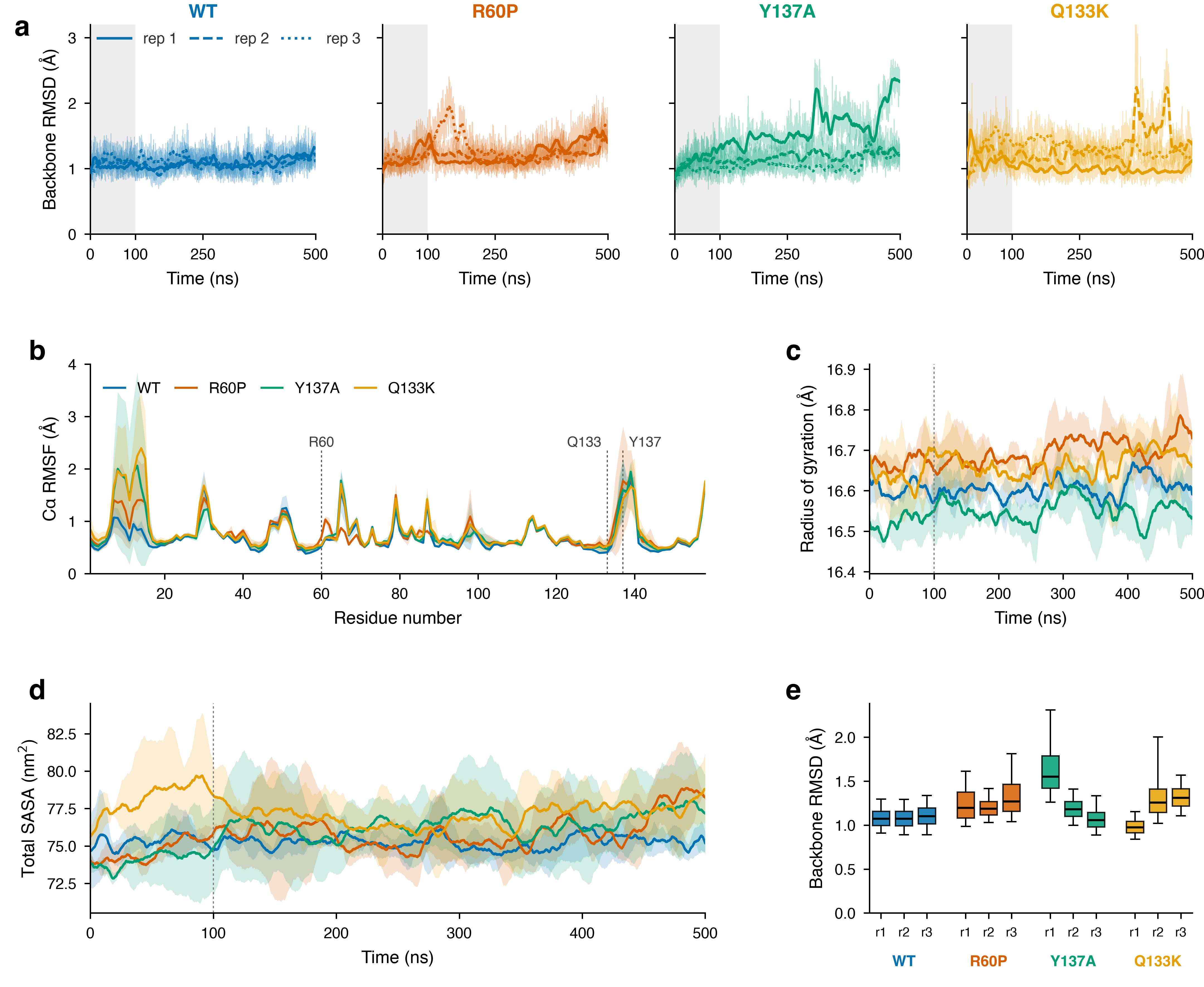

### Figure2_panel_600dpi.png

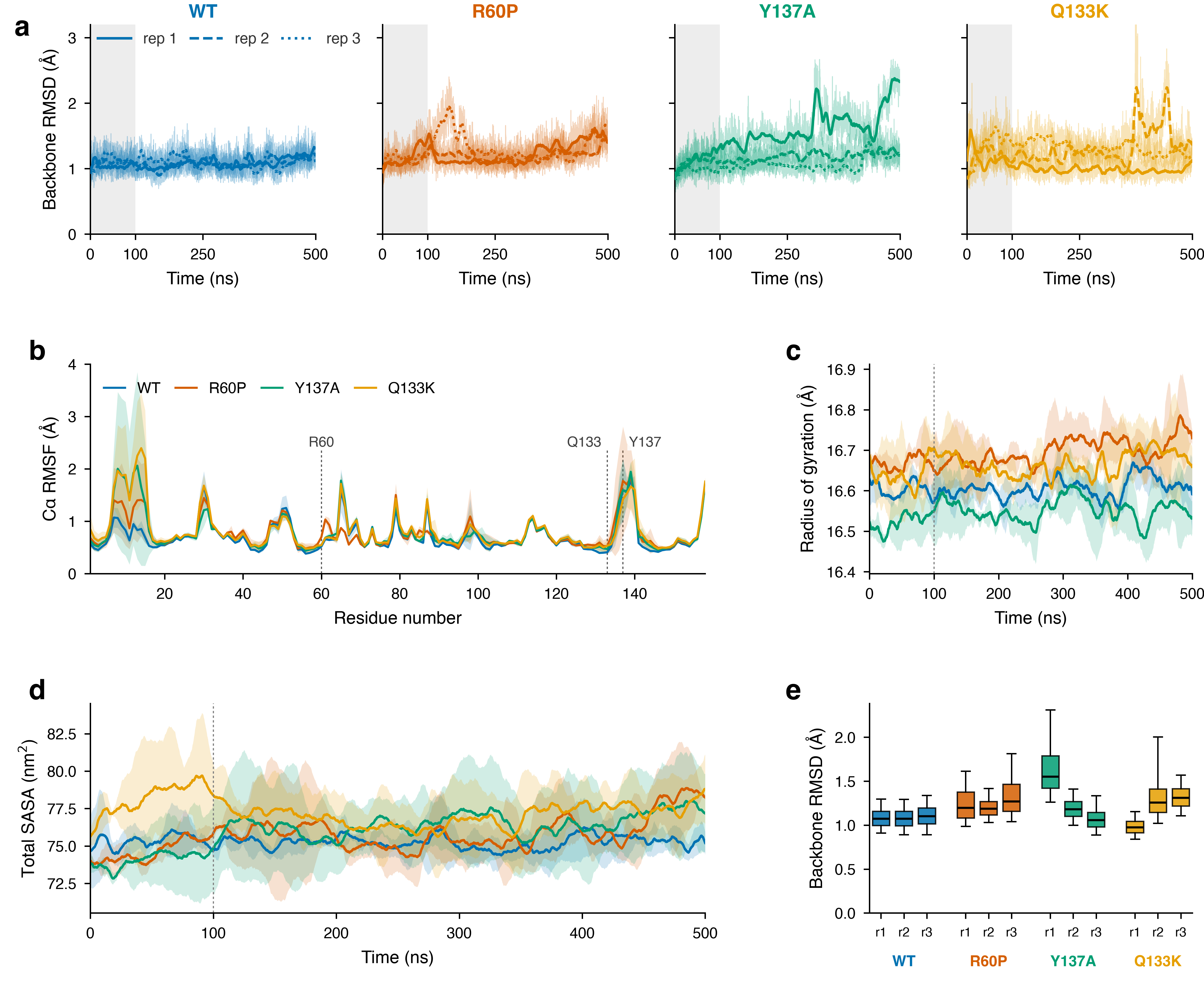

### Figure2_panel_1200dpi.tiff

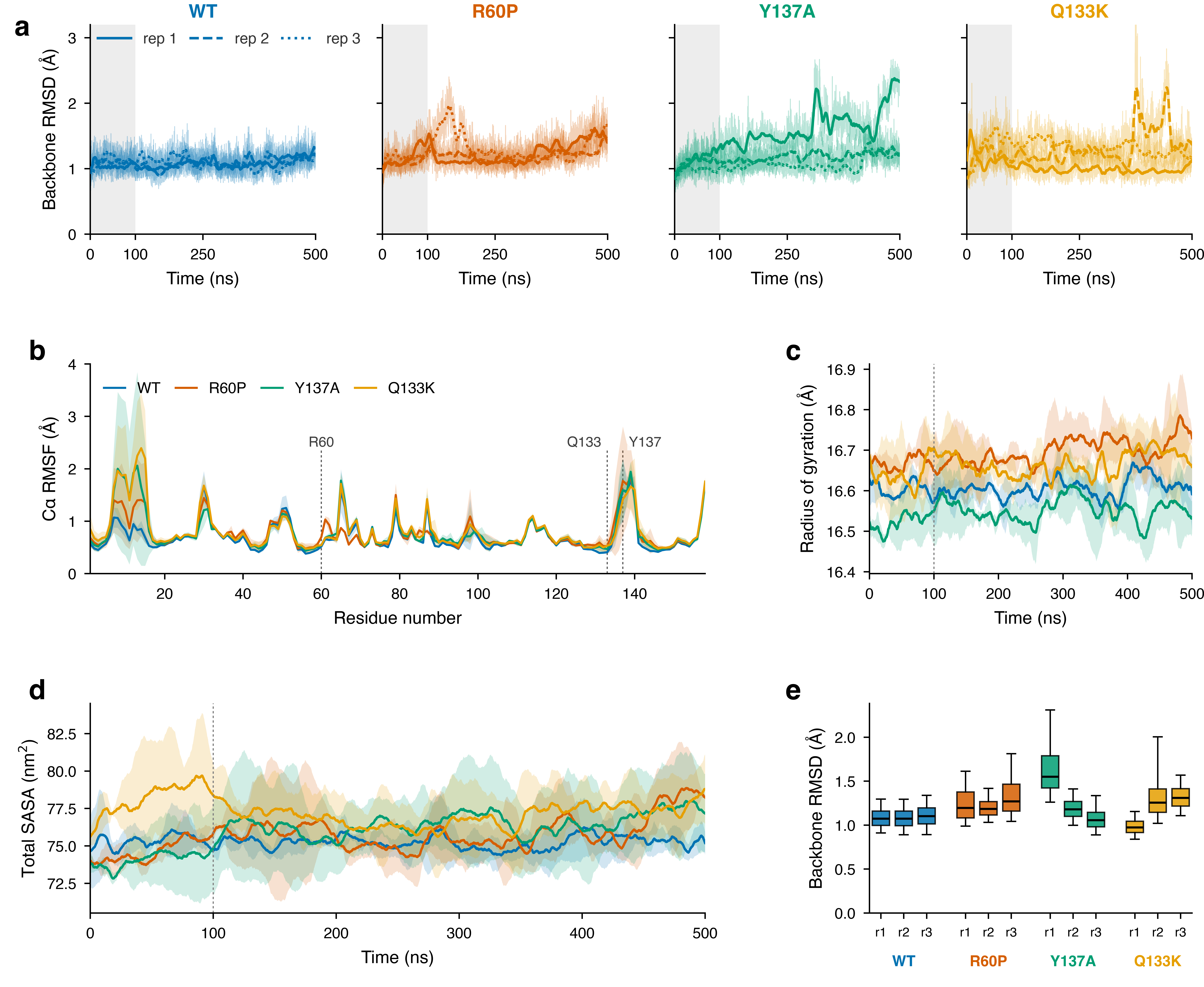

### Figure3_bolsillo_PCA_FEL.jpg

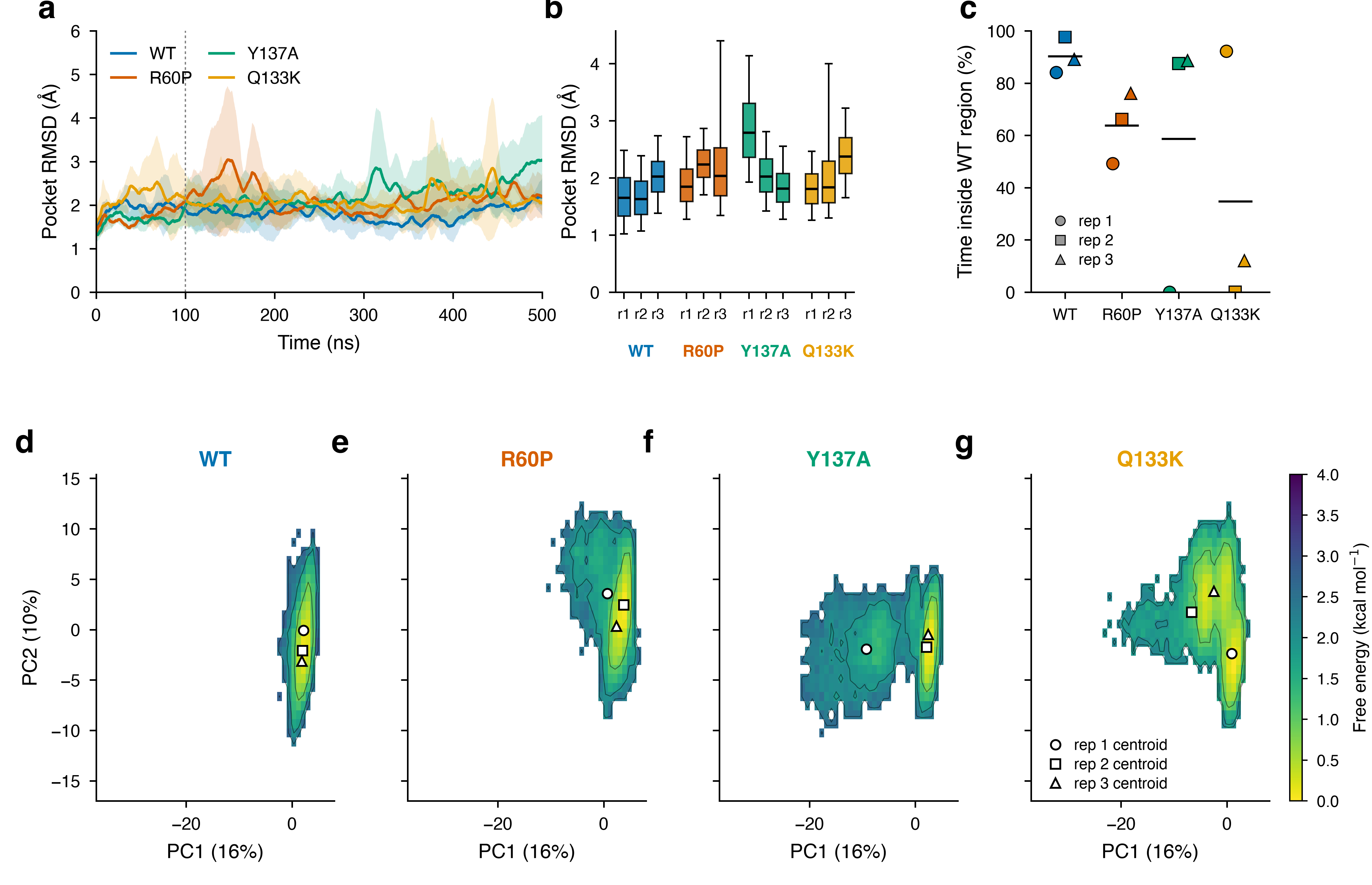

### Figure3_panel_600dpi.png

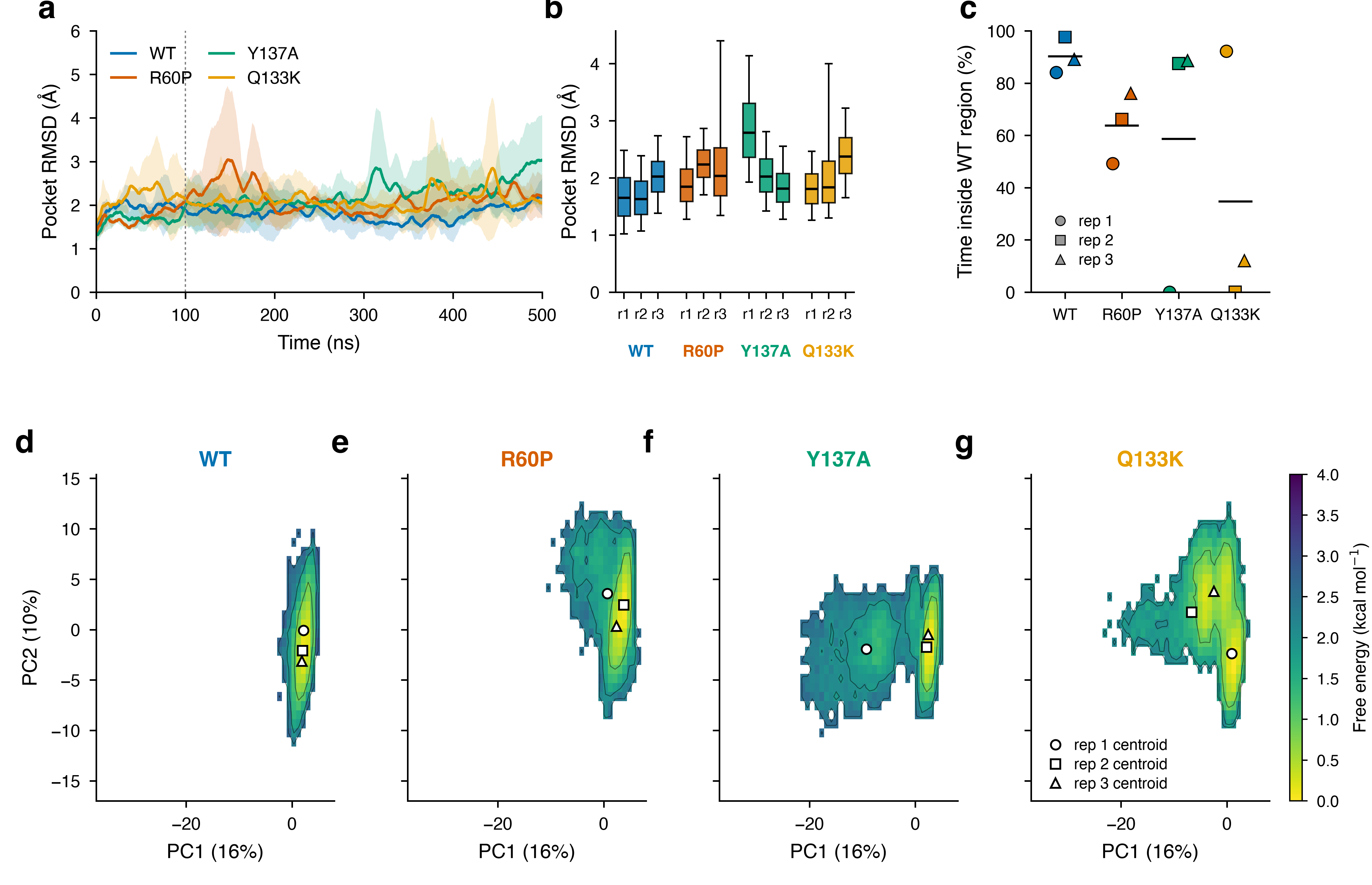

### Figure3_panel_1200dpi.tiff

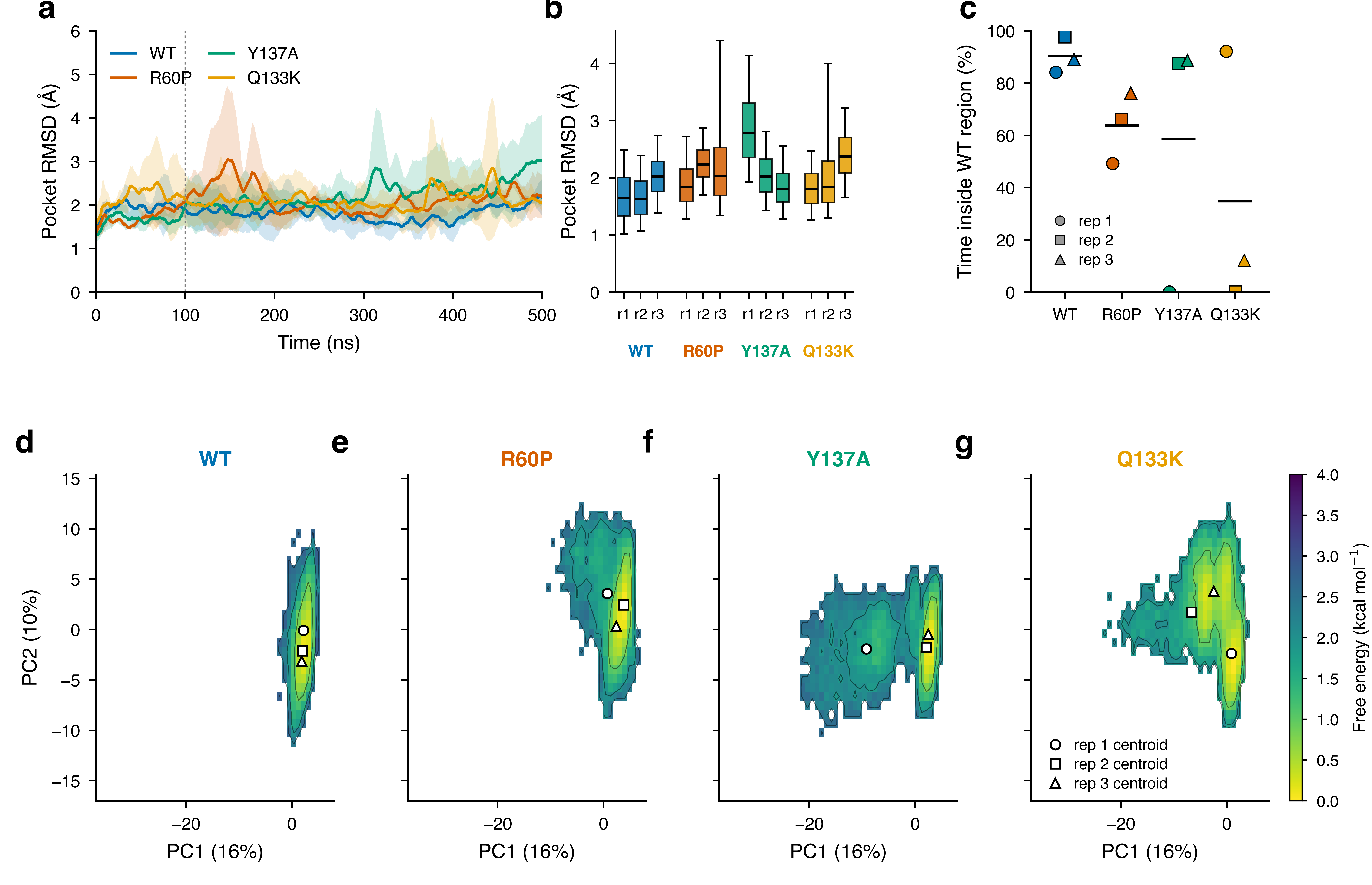

### Figure5_DCCM_estructura_dG.jpg

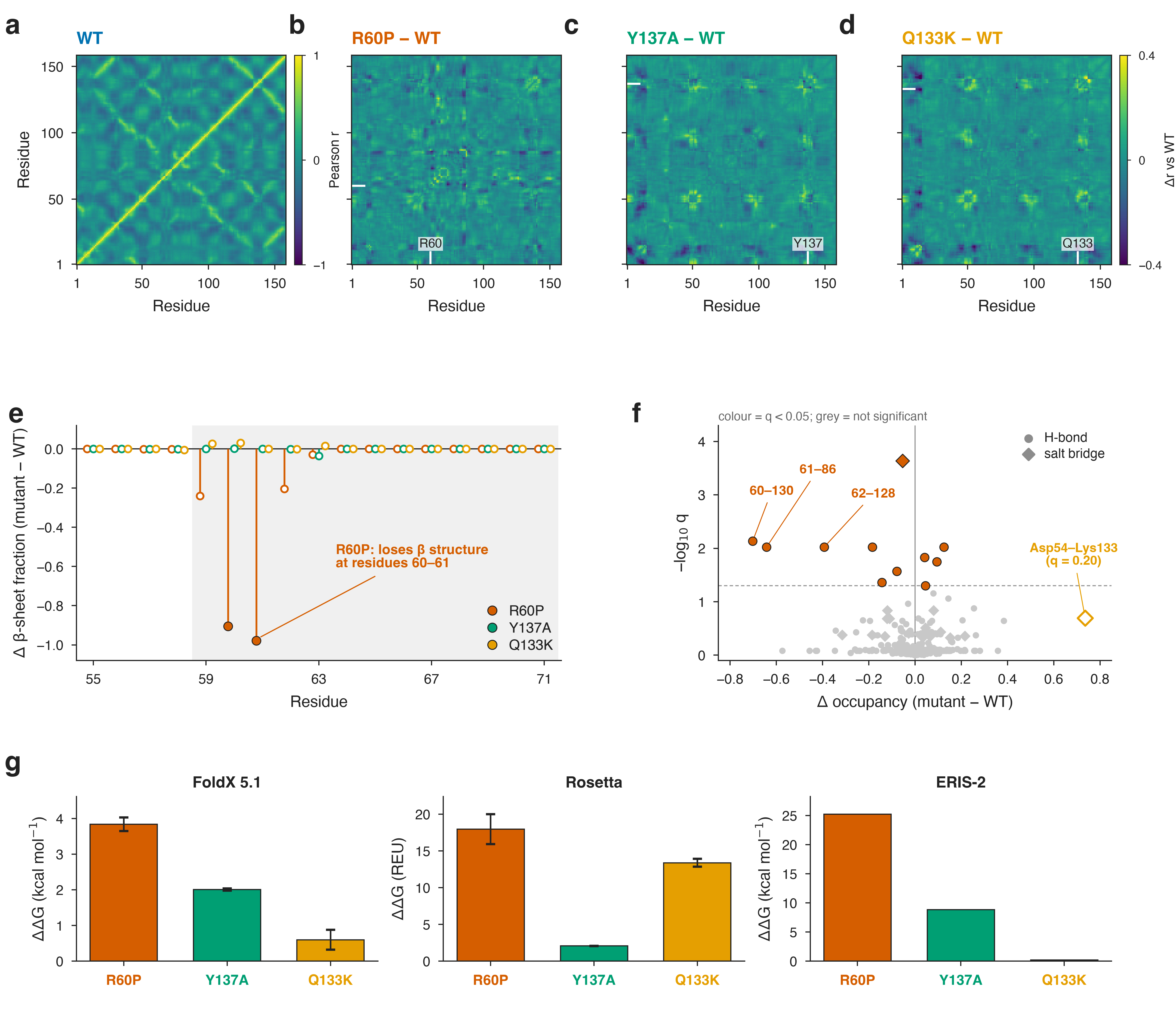

### Figure5_panel_600dpi.png

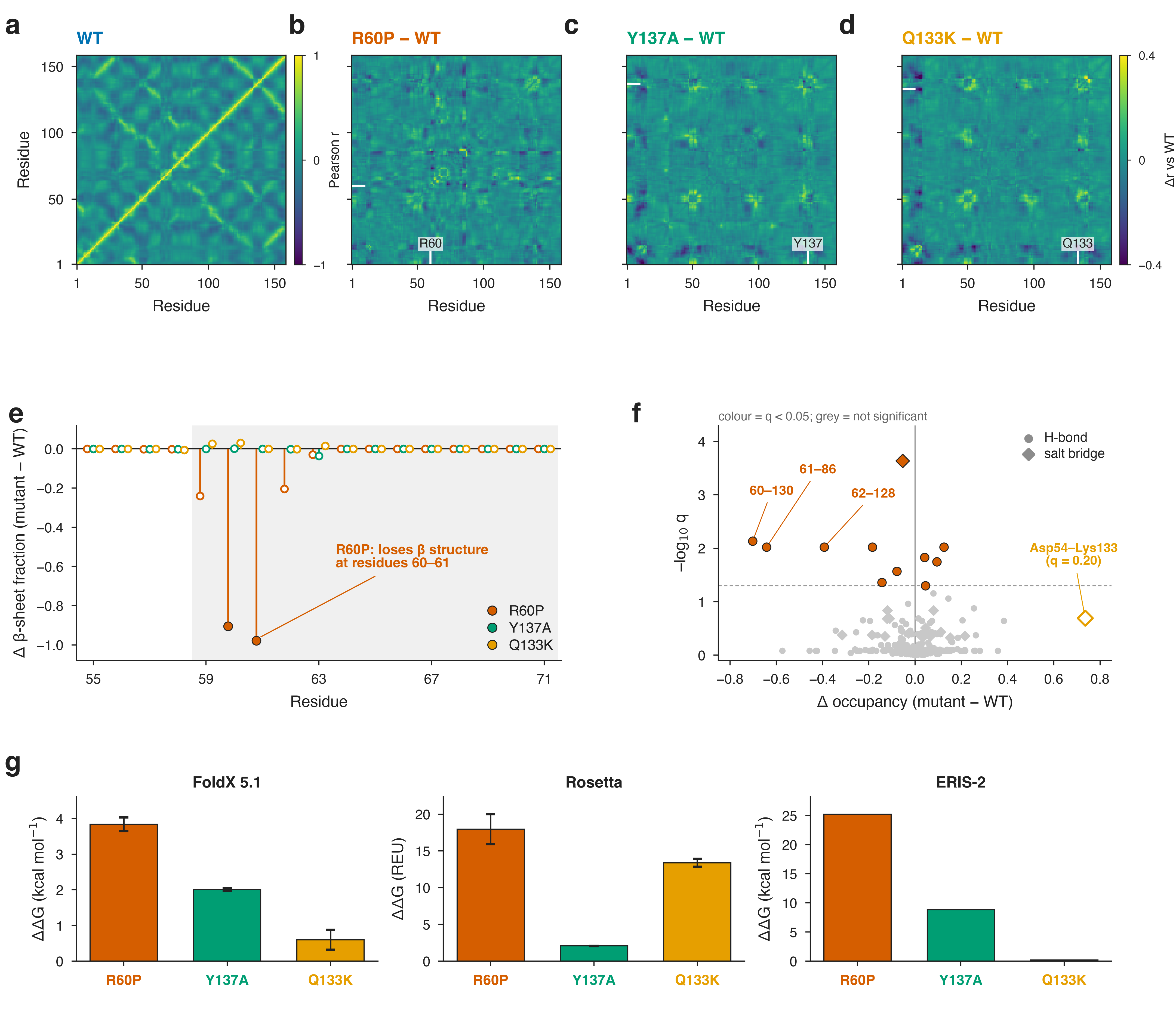

### Figure5_panel_1200dpi.tiff

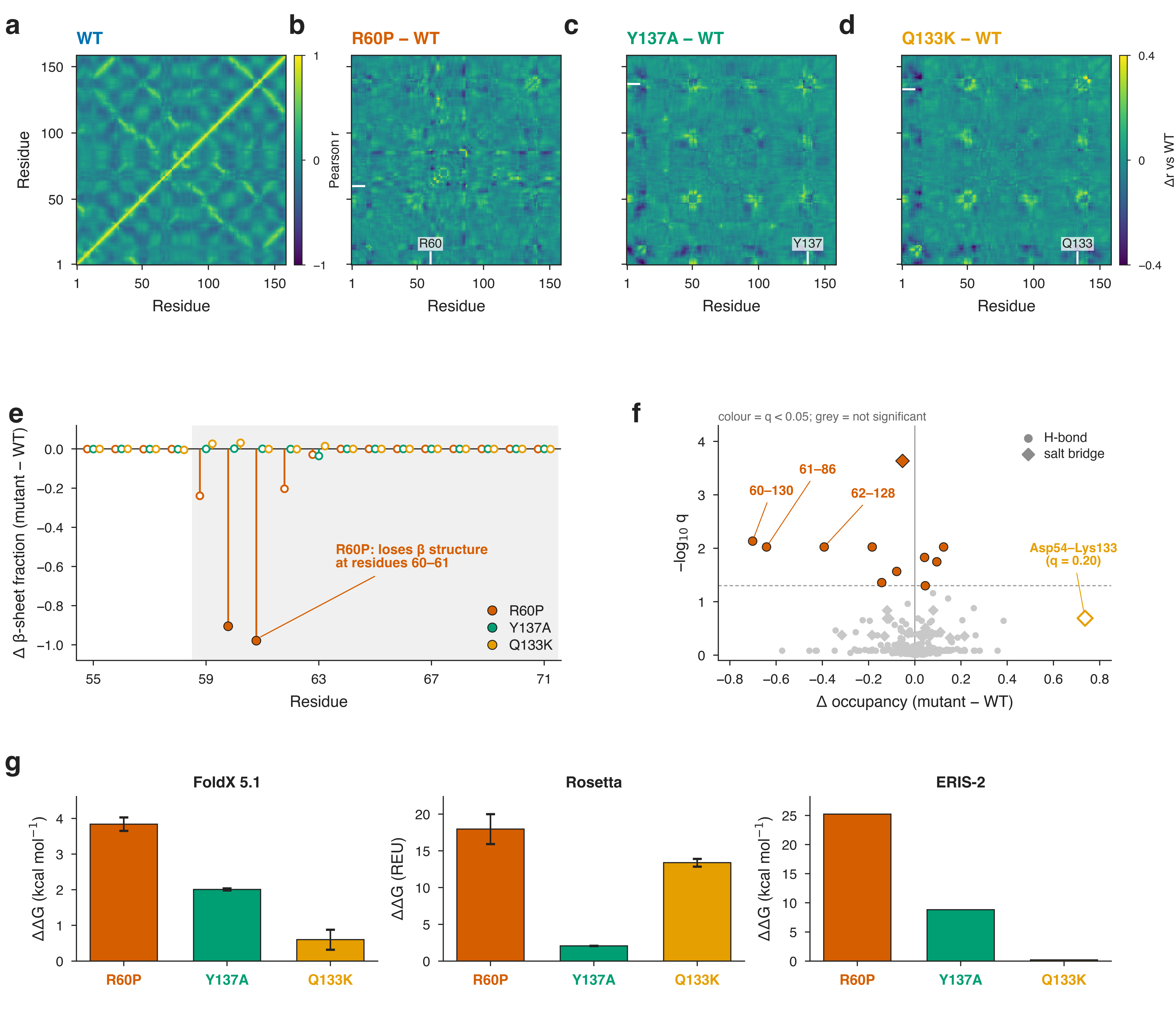

### hires_1.png

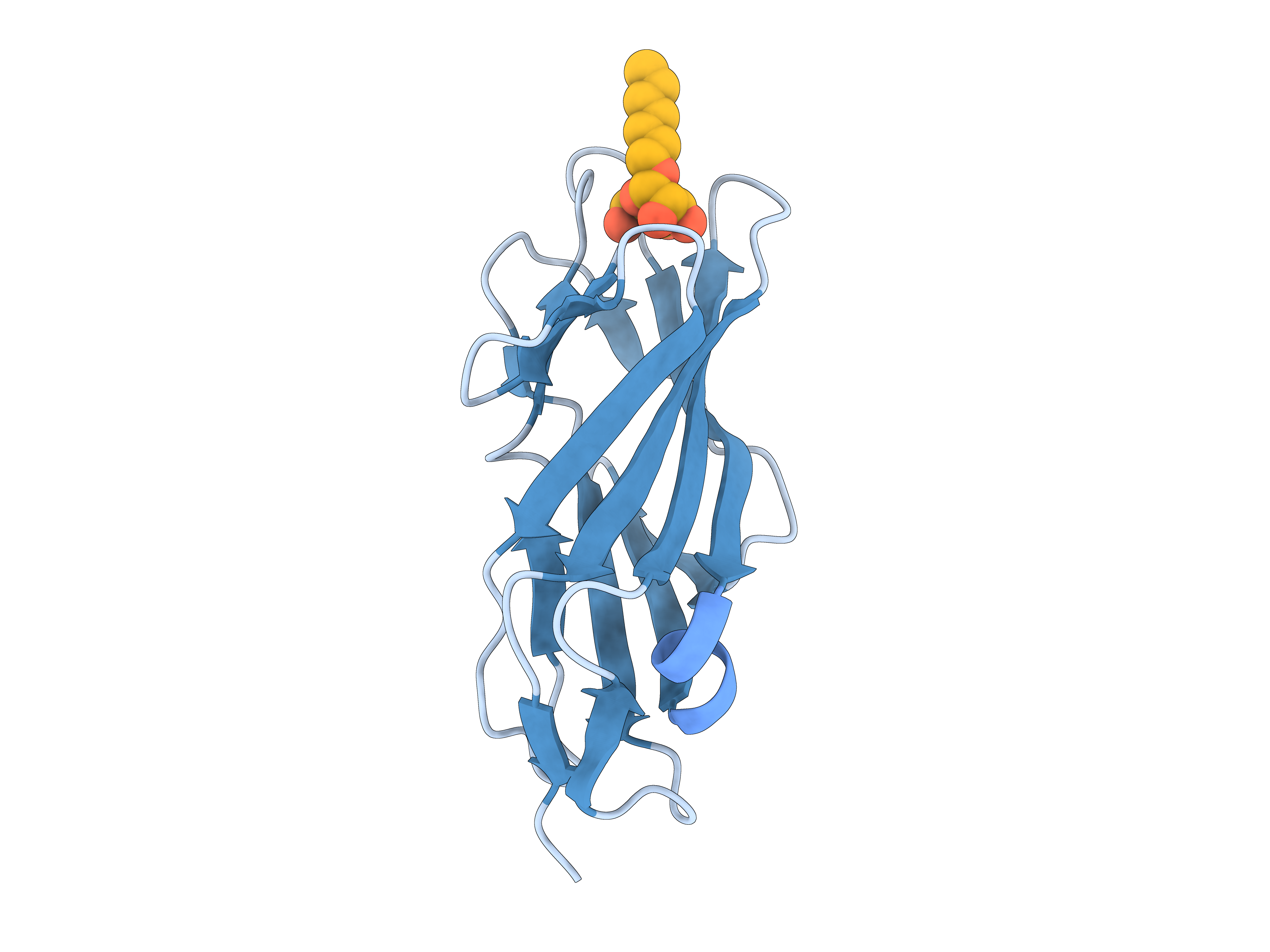

### hires_1_dominios.png

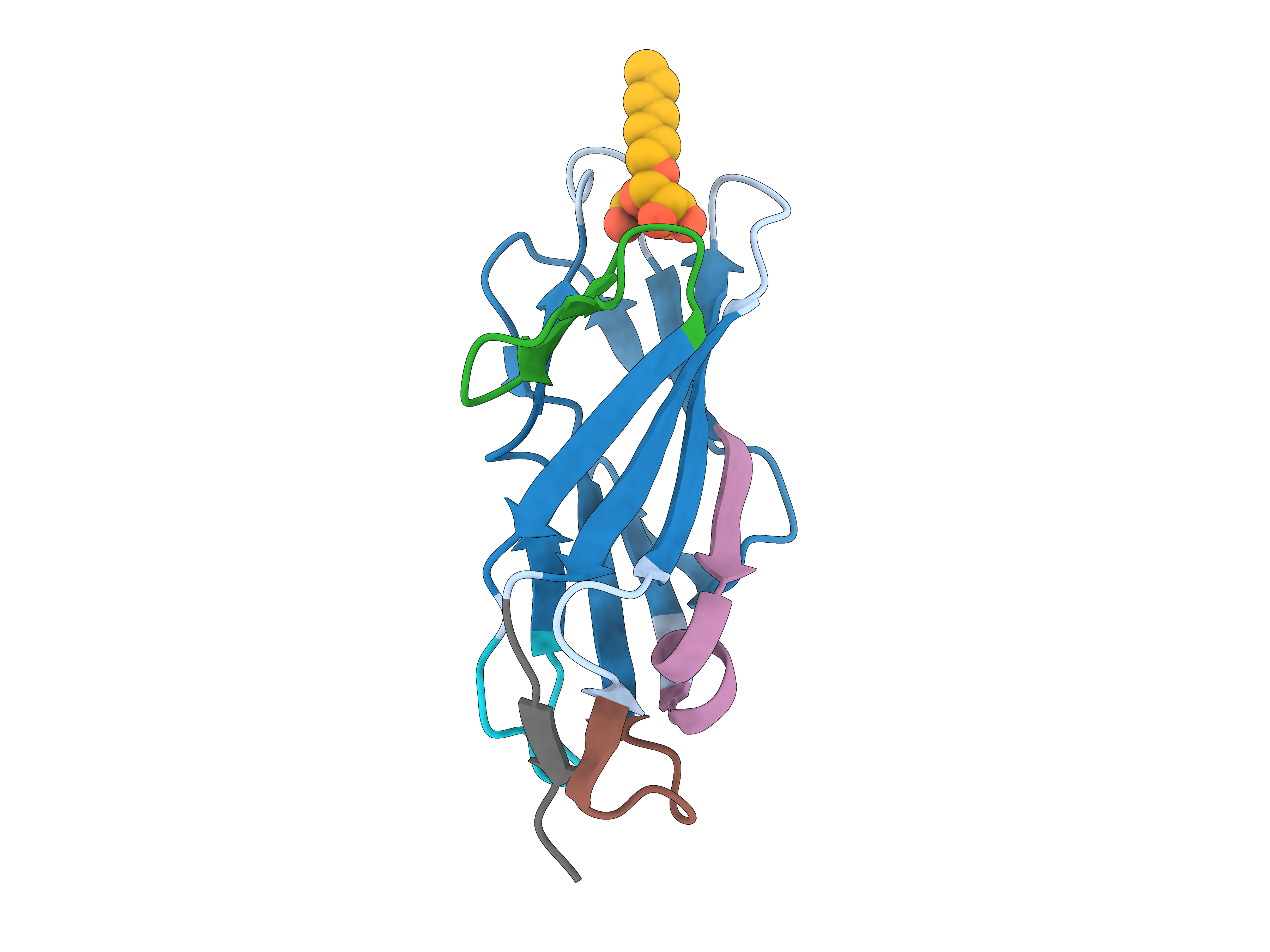

### hires_2a.png

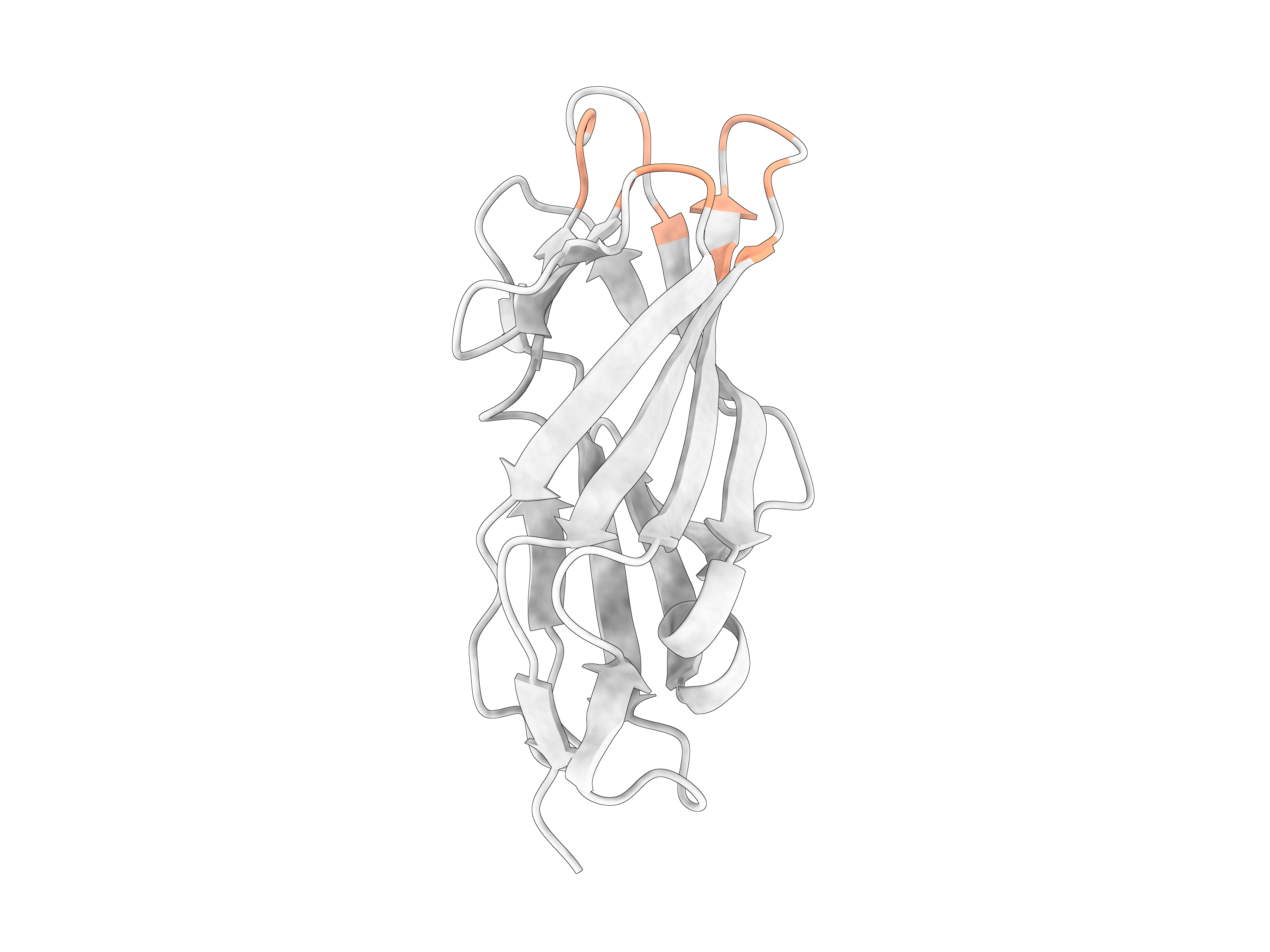

### hires_2b.png

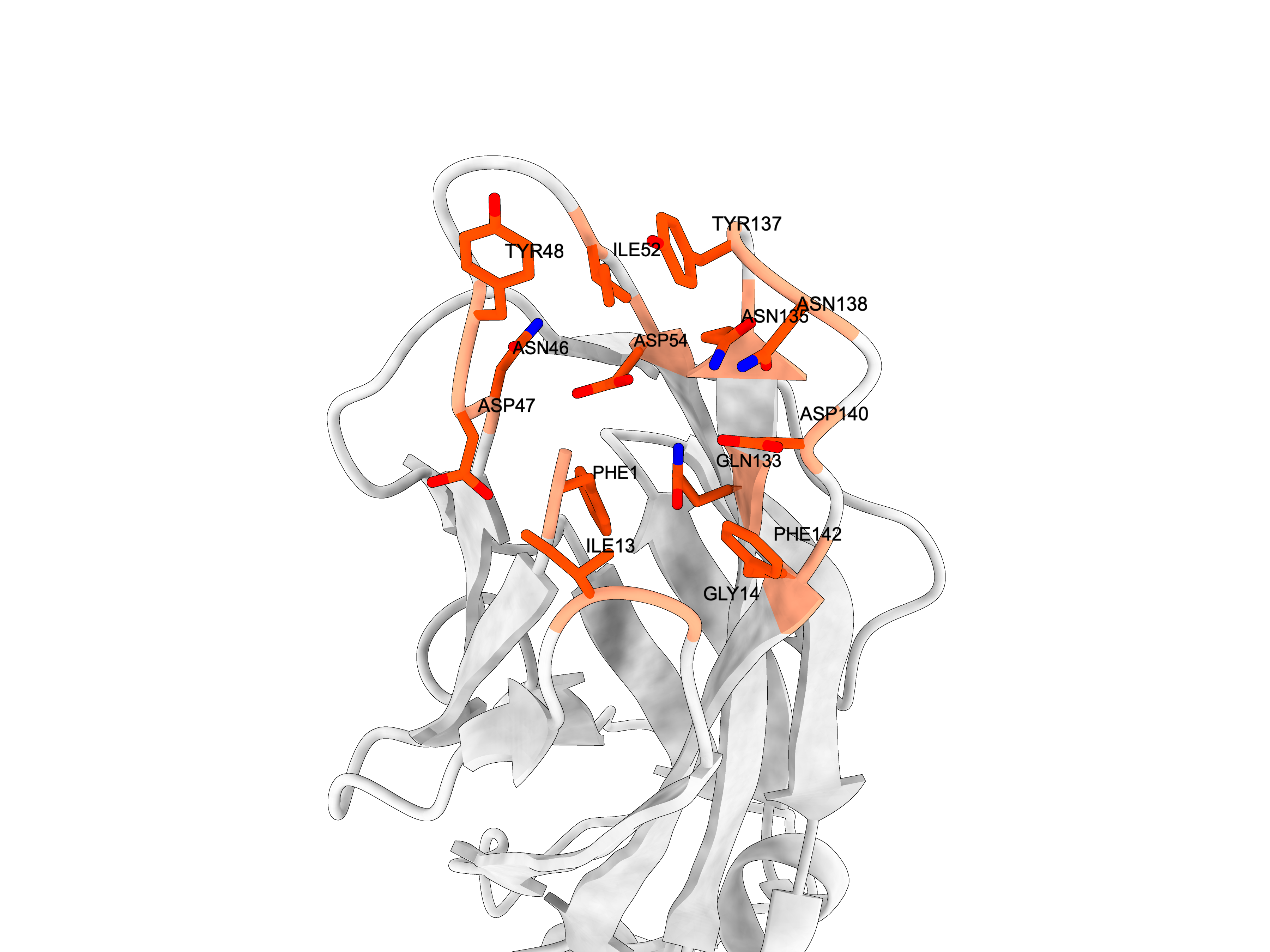

### hires_3.png

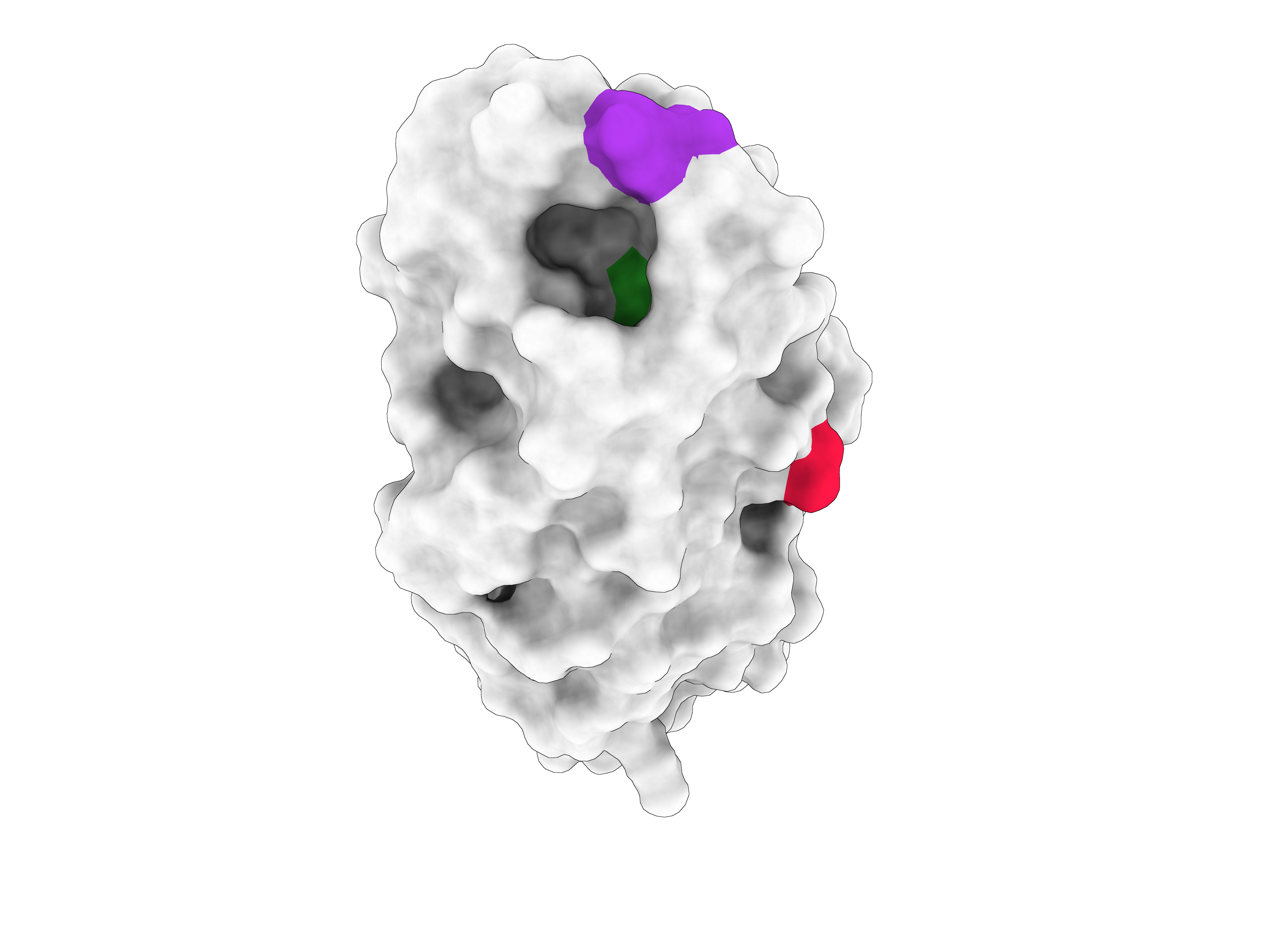

### preview_1_proteina_ligando.png

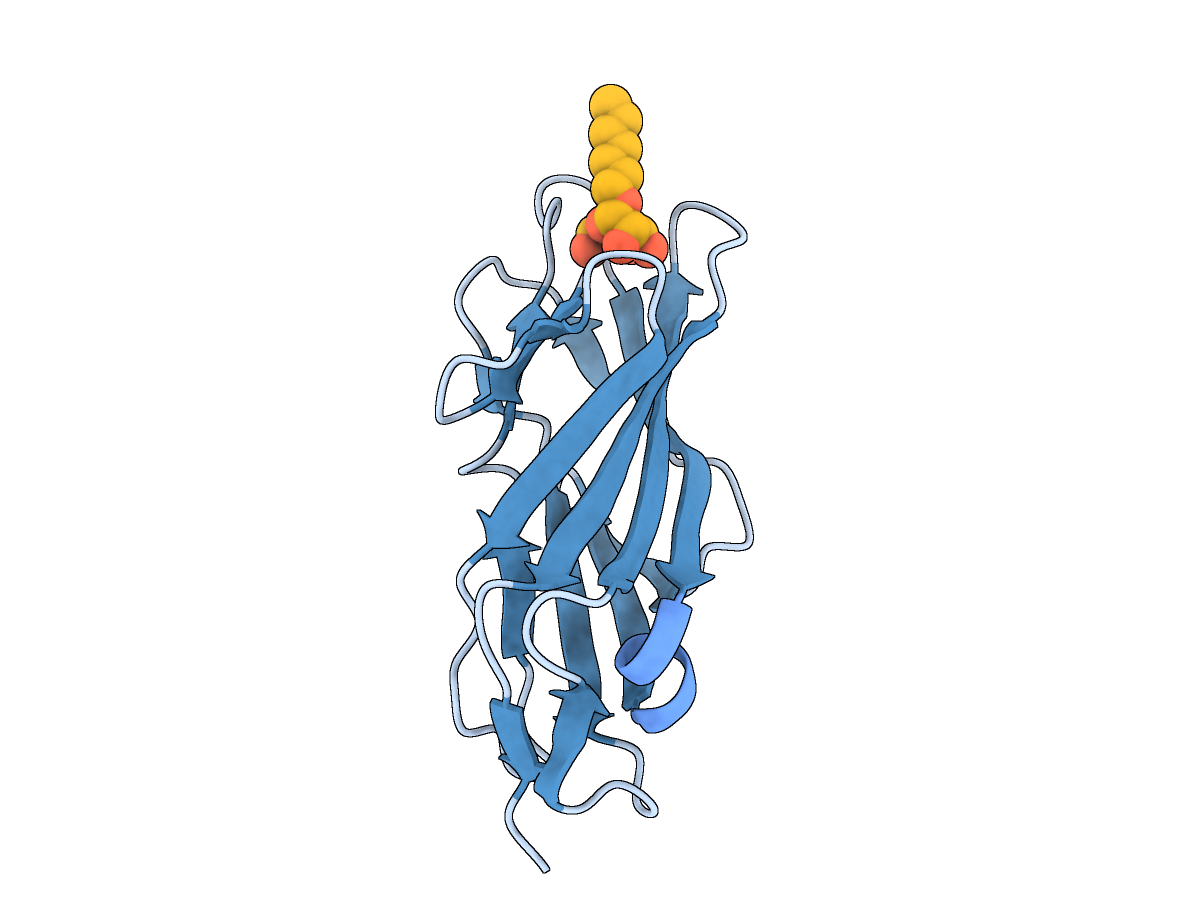

### preview_2_sitio_activo.png

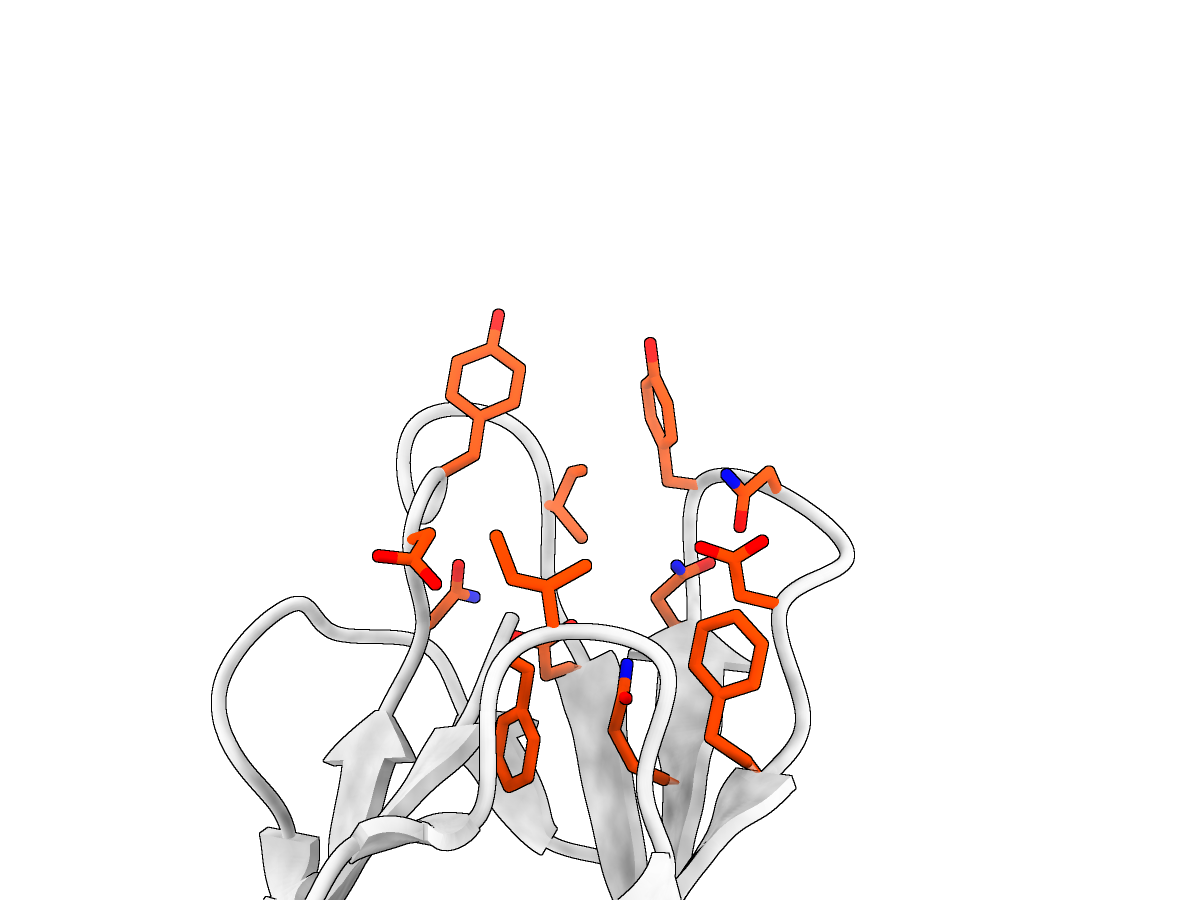

### preview_2a_sitio_activo_general.png

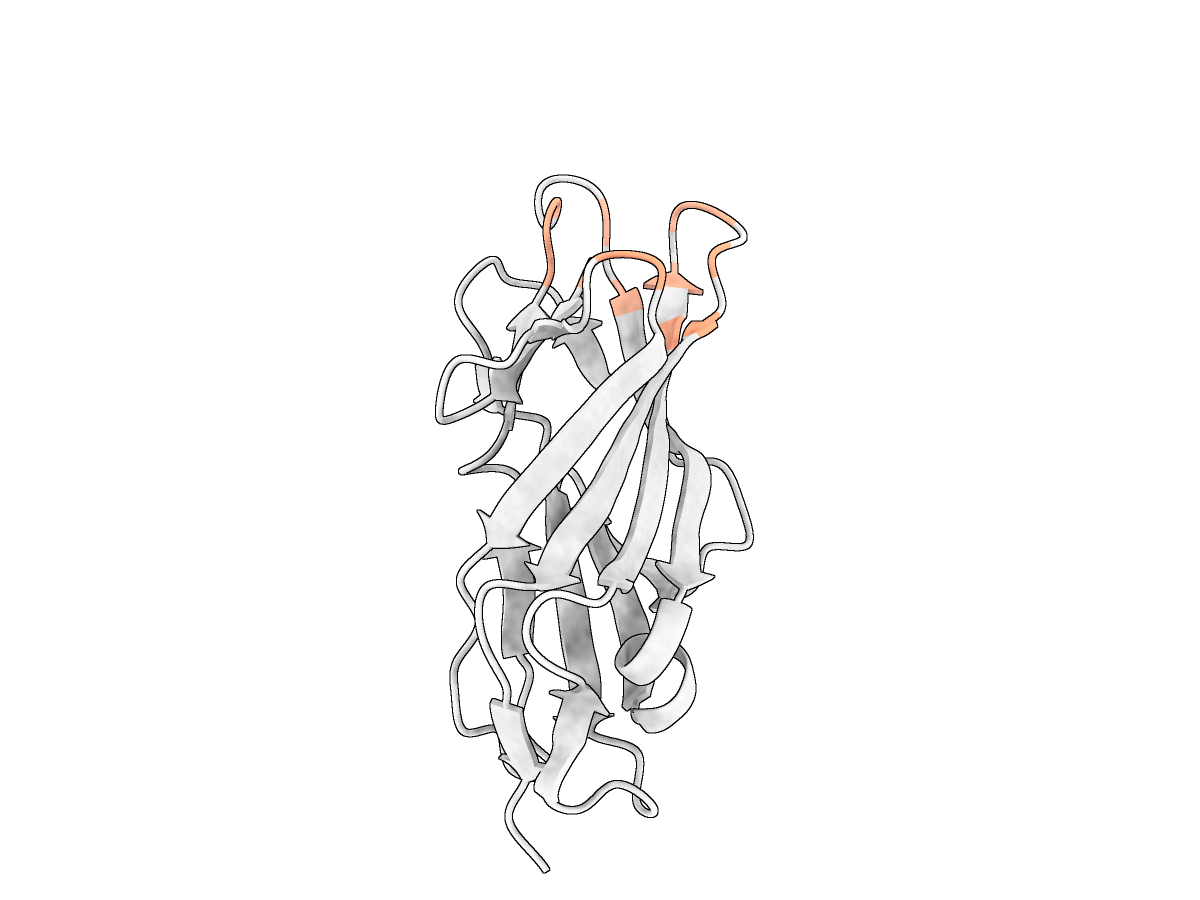

### preview_2b_sitio_activo_cercano.png

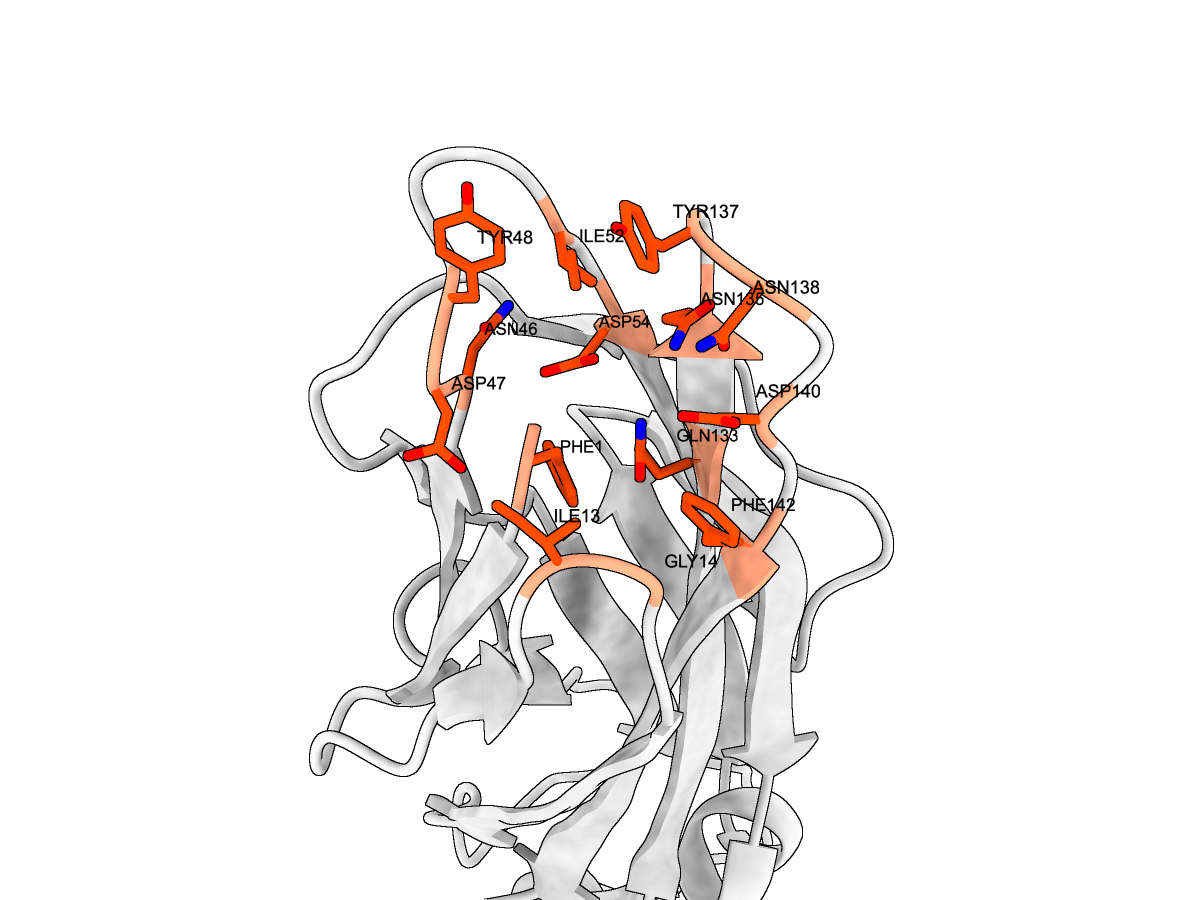

### preview_3_superficie_mutaciones.png

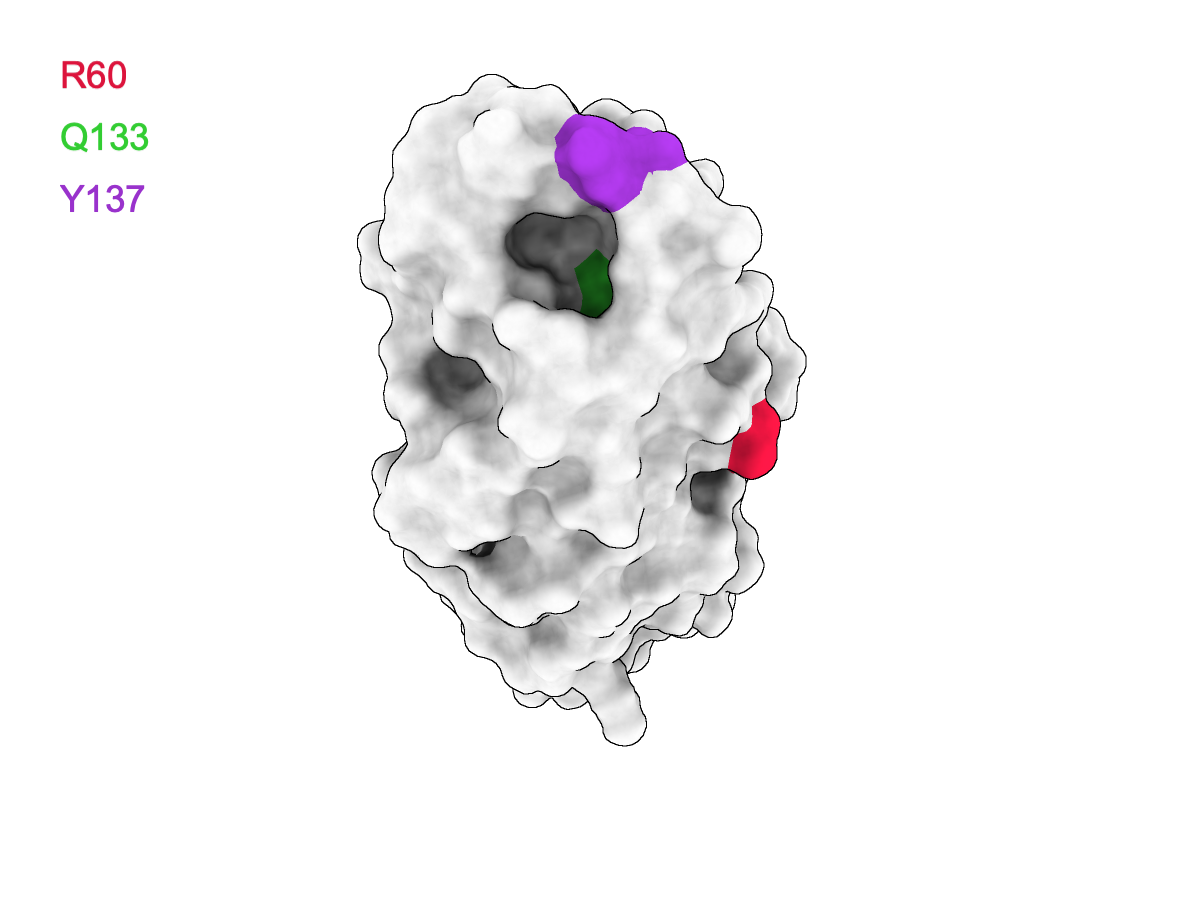
